# OCTOPUS: A versatile open-source tool creating realistic numerical brain cells

**DOI:** 10.64898/2026.09.02.748537

**Authors:** Malte Brammerloh, Inès de Riedmatten, Juliette Beaubis, Jasmine Nguyen-Duc, Rita Oliveira, Andrés Le Boeuf Fló, Elda Fischi-Gomez, Jonathan Rafael-Patiño, Ileana Jelescu

## Abstract

Brain cell morphology plays a crucial role in function and pathology. Biophysical models of diffusion MRI (dMRI) quantify cell morphology *in vivo*, enabling the design of novel biomarkers. These models represent cells by simplified geometries, such as spheres and randomly oriented cylinders, for which analytical signal expressions exist. However, dMRI signals are sensitive to morphological features, such as branching, tapering, undulation, beading, and protrusions, unaccounted for in most models, as they often render analytical solutions impossible. Simulations of the dMRI signal in synthetically generated cells offer a powerful tool to explore how microstructural morphology impacts the dMRI signal. Nevertheless, no tool to generate digital replicas of brain cells is available openly. To address this gap, we introduce the OCTOPUS toolbox, which generates cells featuring all geometrical features described above. OCTOPUS, provided via the Python interface OCTOpool, enables accessible, efficient creation of cells with complex geometries. We recreated histologically reconstructed neuronal and glial cells, including pyramidal, GABAergic, and glutamatergic neurons, and astroand microglia. To illustrate the plausibility of OCTOPUS-generated cells, we reproduced established properties of dMRI signals from brain tissue, such as the signatures of short-range disorder, branching and protrusions, and a high-b-value power law. By comparing the geometries and dMRI signals of generated and original cells, we found different growth strategies adequate for more isotropic and more anisotropic cells. We anticipate that realistic cell substrates created by OCTOPUS will help validate biophysical models, design dMRI sequences sensitive to fine-grained cell morphology beyond analytical models, and generate realistic numerical substrates of brain tissue.

**Highlights:**

- OCTOPUS is an open-source tool generating realistic gray matter cells in seconds.
- Cells have tunable features: branching, tapering, undulation, beading, and spines.
- A Python wrapper facilitates fast, flexible cell replication with OCTOPUS.
- dMRI decays in cells showed signatures of disorder, branching, and spines.
- OCTOPUS is a powerful tool to explore microstructure imaging techniques.

## 1. Introduction

Since the seminal work of Ramón y Cajal, histological methods have enabled unparalleled insight into brain cell morphologies and their pathological alterations. Nonetheless, they are restricted to *post mortem* tissue and are further limited by their time-consuming and costly nature. Thus, while histology serves as the gold standard for defining pathologies, its clinical applicability is limited, underscoring the need to develop *in vivo* histology techniques (Weiskopf et al., 2021).

To meet this need, diffusion magnetic resonance imaging (dMRI) quantifies tissue microstructure *in vivo* (Jelescu et al., 2020). It leverages sensitivity to water displacement to probe the underlying microstructure within the imaging voxels. As water molecules encounter obstacles such as cellular membranes, their surrounding microstructural environment restricts intra- and extracellular water diffusion. Similarly, diffusion-weighted magnetic resonance spectroscopy (dMRS) is sensitive to the diffusion of intracellular metabolites (Ligneul et al., 2024). As different metabolites are preferentially found in distinct cell types, such as neurons and glial cells, they act as cell-specific microstructure probes.

Informed by dMRI and dMRS signals, analytical biophysical models were developed to estimate specific tissue parameters, particularly in healthy white matter and gray matter (Stanisz et al., 1997; Zhang et al., 2012; Jelescu et al., 2022; Palombo et al., 2020; Novikov et al., 2019; Alexander et al., 2019), and in tumors (Panagiotaki et al., 2014; Voronova et al., 2025). They approximate tissue microstructure using simple geometries, such as spheres, sticks, and cylinders for which analytical closed-form solutions exist. By fitting these analytical models, parameters such as soma and cell process fractions, soma and cell process radii (particularly axon radii), membrane exchange times, cell process orientations, and intra- and extracellular diffusivity can be estimated. However, geometrical simplifications assumed by biophysical models do not capture more fine-grained features of cell morphology. Several studies examined how the dMRI signal and biophysical models thereof are impacted by morphological features, among them branching (Palombo et al., 2016), tapering, undulation and beading (Lee et al., 2020a; NguyenDuc et al., 2026b; Oliveira et al., 2026), and protrusions (Palombo et al., 2017; Şimşek et al., 2025; Chakwizira et al., 2025), including dendritic spines and astrocytic leaflets. These studies have examined these features separately, often using simplified cell representations (Şimşek et al., 2025; Chakwizira et al., 2025; Palombo et al., 2016; Nguyen-Duc et al., 2026b; Oliveira et al., 2026). Assessing their combined and relative contributions in realistic cells, therefore, requires a unified framework that jointly represents these features. Such a framework is currently lacking. Hence, while biophysical models based on dMRI and dMRS are promising tools for *in vivo* histology, they require further validation to determine their range of validity.

Numerical simulations mimicking diffusion in tissue are a flexible tool for studying the dMRI signal in realistic yet controlled settings, enabling validation of biophysical models and informing data-driven models for advanced microstructure characterization. To simulate the dMRI signal, an accurate digitized representation of brain tissue is pivotal. To generate these substrates, two main approaches are employed. A substrate can be obtained from histological data, for instance, by segmenting light (Tecuatl et al., 2024), proton-beam (Brammerloh et al., 2021), or electron (Lee et al., 2020a) microscopy. Although histological substrates rely on gold-standard techniques for imaging cellular morphologies, their accuracy is limited by tissue alterations during fixation, segmentation inaccuracies, and imperfect 3D reconstruction. Moreover, high-quality histological cell reconstructions are scarce, as their precise segmentation requires extensive work (Augustin et al., 2025). Therefore, synthetic tissue generators offer a powerful alternative to study the relationship between complex microstructural features and the dMRI signal. Such generators allow freedom over the generated cell morphologies, enabling the representation of geometric alterations due to pathology (Mosso et al., 2024; Budde and Frank, 2010), brain development (Ligneul et al., 2025), and functional activation (Spencer et al., 2025). They also enable the creation of a large cell dictionary with controlled, gradual parameter changes; more comprehensive than achieved using histological segmentation.

Tissue generators are largely described and used for substrates made of densely packed axons, such as those found in white matter (Ginsburger et al., 2019; Villarreal-Haro et al., 2023; Callaghan et al., 2020; Winther et al., 2024; Nguyen-Duc et al., 2026a). Regarding gray matter cells, efforts to develop cell generators have been underway in recent years. The first comprehensive single-cell generator was reported by Palombo et al. (2019), although it was not released as open-source. Then, MEDUSA (Ginsburger et al., 2019) and CATERPillar (Nguyen-Duc et al., 2026a) focused on modeling astrocyte morphology in white matter, without supporting other gray matter cell types. More recently, the first generator of a realistic cortical column was presented (Aird-Rossiter et al., 2026b). Currently, however, no openly accessible gray matter cell generator is available that allows for freely tuning the cellular geometry.

A fundamental requirement for any cell generator is the ability to produce biologically plausible cells. This can be validated by directly quantifying geometric parameters of these cells, such as their extent, branching order, and process radius. Indirectly, the cell’s plausibility can be corroborated by reproducing well-established dMRI signal signatures in biological substrates. For instance, it has been demonstrated that cell process complexity, such as beading, falls within the short-range disorder class (Novikov et al., 2014; Lee et al., 2020b), which implies the absence of long-range correlations. Short-range disorder is reflected in the diffusion time dependence of the dMRI signal: both diffusivity and kurtosis show a time dependence following 1/√*t* (Novikov et al., 2014; Lee et al., 2020b). Furthermore, past research found that the dMRI signal from white matter follows a power law of 1/√*b* at high b-values in the long-diffusion-time limit (Veraart et al., 2019). This power law is expected for randomly oriented sticks, i.e., impermeable, one-dimensional objects (Callaghan et al., 1979). In simulations of gray matter cells, this power law holds for elongated impermeable cell processes of sufficiently small radius, while the cell soma has an additional contribution (Olesen et al., 2022). Moreover, exchange between different cellular branches decreases the apparent diffusion coefficient at long diffusion times, enabling the quantification of branch lengths using dMRS (Palombo et al., 2016). Recently, the effect of dendritic spines on the dMRI signal has been examined, and a reduction of the effective diffusivity has been reported (Şimşek et al., 2025; Chakwizira et al., 2025). Hence, dMRI signals simulated in realistic cells should have these properties.

This study presents the OCTOPUS toolbox, an open-source realistic brain cell generator. In contrast to most substrates, which are defined with triangular meshes, we represent the cells in a memory-efficient manner using overlapping spheres, as introduced in (Ginsburger et al., 2019) and later adopted in (Nguyen-Duc et al., 2026a). First, we describe the implementation details: how characteristic morphologies of different cell types are obtained from histological cell reconstructions and enforced in the synthetic cell generator. In particular, beading and undulation are newly modeled by stochastic processes. We also provide a brief overview of the OCTOpool Python wrapper of OCTOPUS. Next, we validate the biological plausibility of OCTOPUS-generated microstructure by examining whether dMRI signals are consistent with short-range disorder and the stick power law. We demonstrate the flexibility of OCTOPUS by replicating a broad range of histological cell reconstructions, including pyramidal, GABAergic, and glutamatergic neurons, as well as several glial cells, using different growth strategies. The generated cells are compared to the original cells segmented from histology by evaluating the parameters obtained using OCTOpool, their morphology using Sholl analysis, and their dMRI signals. Eventually, we examine the signatures of branching and spines in the dMRI signal.

## 2. Methods

The core methodology of this paper is the OCTOPUS toolbox, which we describe in the first subsection. Next, we outline the OCTOpool toolbox, a Python wrapper for using OCTOPUS to generate cell replicas, run dMRI Monte Carlo simulations, and analyze their results. Lastly, we describe the virtual experiments we performed to validate the OCTOPUS toolbox.

### 2.1. OCTOPUS

The OCTOPUS toolbox enables creating synthetic cells with tunable morphological complexity, such as branching, tapering, undulation, beading, and protrusions (Figure 1A). It generates cells by placing a soma, to which it adds primary branches. From the endpoints of the primary branches, secondary processes are grown. Subsequently, more and more secondary processes are grown iteratively on all available branch endpoints. Eventually, protrusions, such as spines, grow on the branches. Cell growth is controlled by an OCTOPUS configuration file, whose parameters are listed in Table 1. These input parameters can be either manually input, for instance following tables of histological data (Aird-Rossiter et al., 2026a), or directly extracted from 3D histological cell segmentations, for instance obtained from (Ascoli et al., 2007), using the OCTOpool toolbox (Figure 1B, see subsection 2.2). In the following, we describe the growth algorithm and all the configuration parameters in detail. More information can be found in the repository, which will be released upon publication of this manuscript.

**Figure 1:**
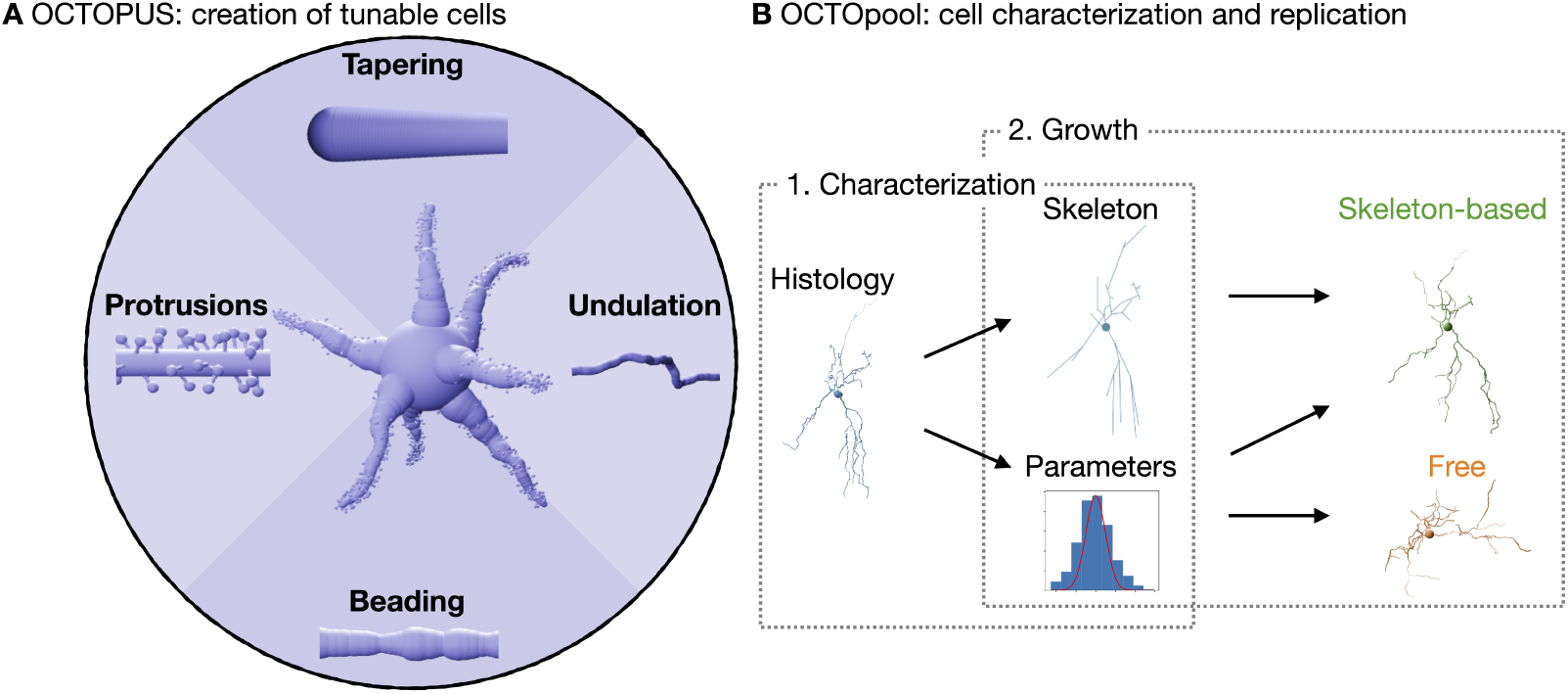
Overview of the features of OCTOPUS and OCTOpool. A) An example cell exhibiting tapering, undulation, beading, and protrusions illustrates the features included in the OCTOPUS toolbox. Each morphological feature is illustrated within separate quadrants of the disk. B) The OCTOpool toolbox enables: 1. characterization of morphological parameters (such as branch radii, branching order, etc.) and extraction of the cell skeleton from histological 3D cell reconstructions; 2. growth of cells using OCTOPUS with a free or skeleton-based approach. The free approach generates the cell architecture from input parameters, whereas the skeleton-based approach preserves the cell branching structure and main branch directions of the histological reconstruction. Controlled features, such as beading, undulation, and spines, can be added using either approach.

**Table 1.**
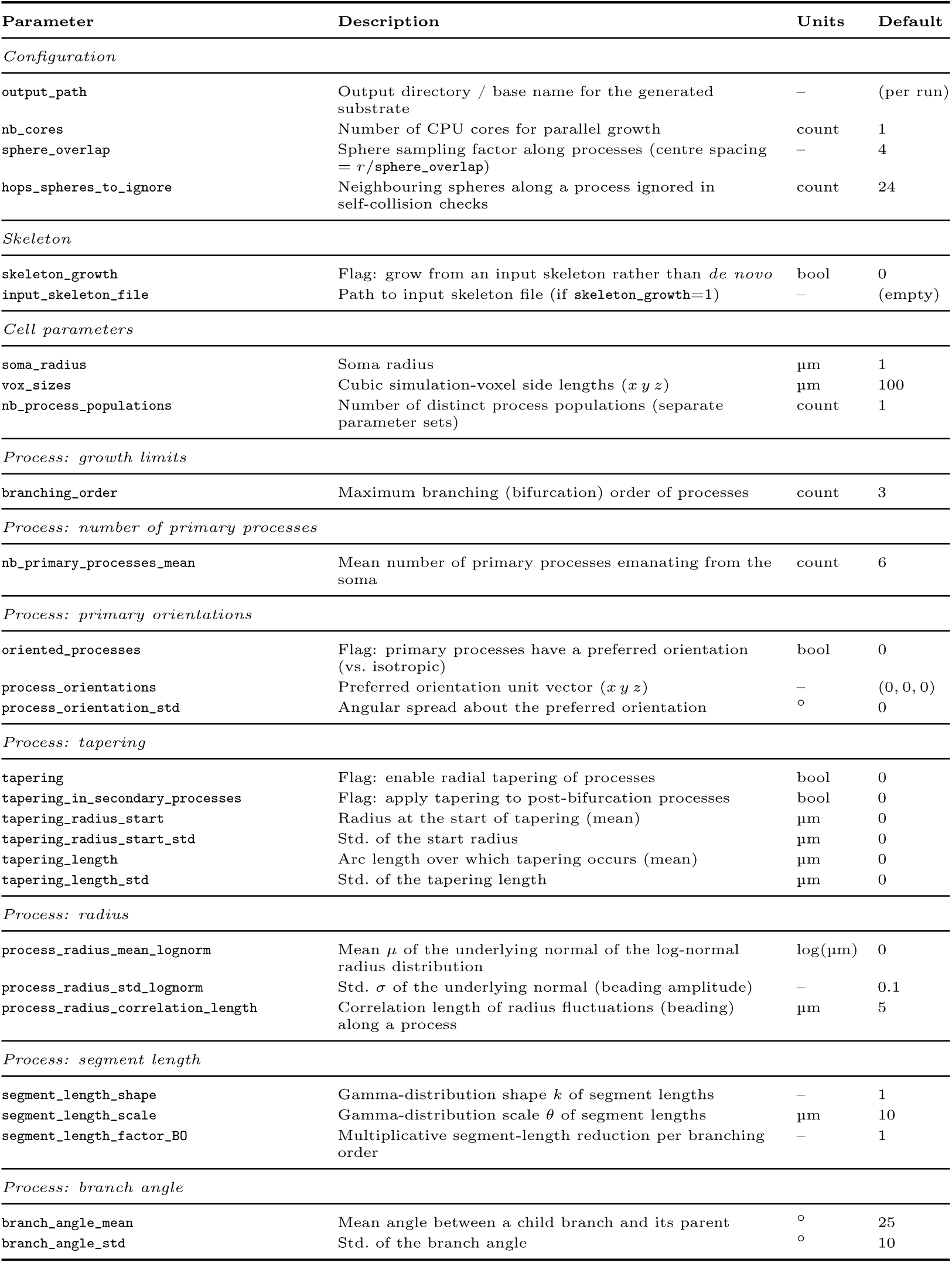
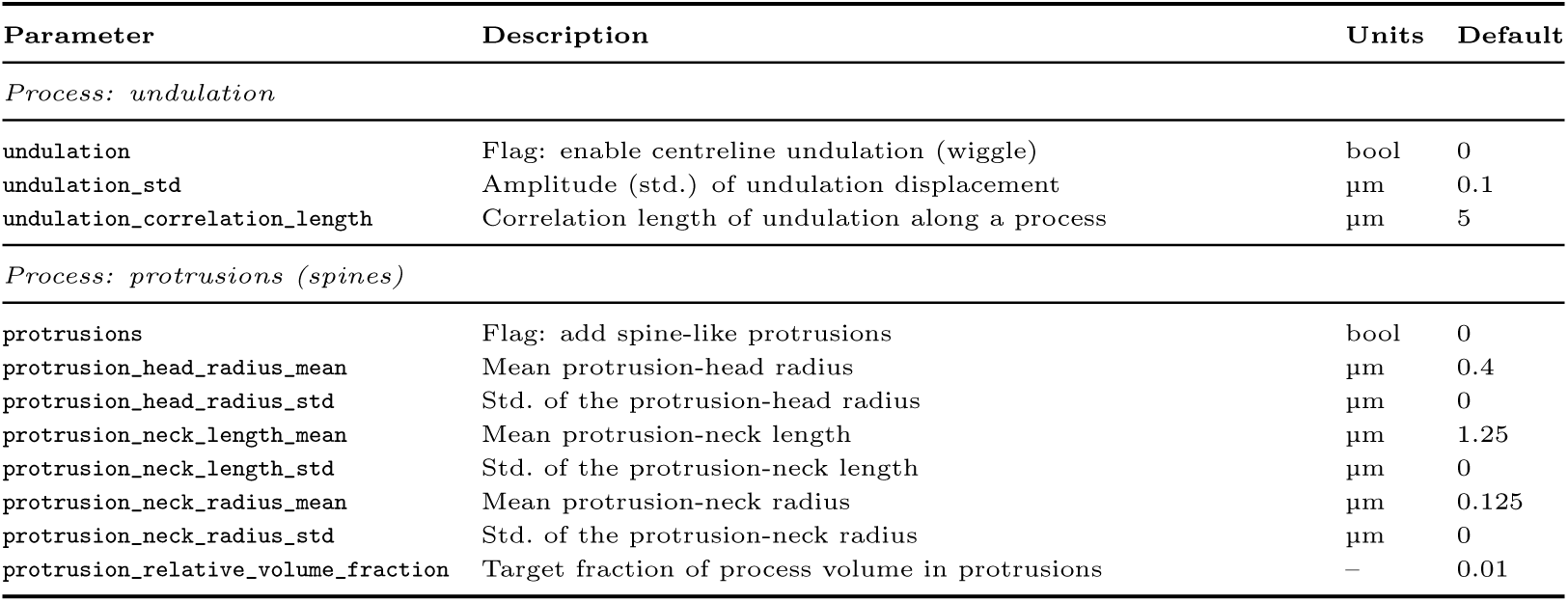
OCTOPUS cell-generator input parameters, with descriptions, units, and default values.

| Parameter | Description | Units | Default |
| --- | --- | --- | --- |
| *Configuration* |  |  |  |
| output_path | Output directory / base name for the generated substrate | – | (per run) |
| nb_cores | Number of CPU cores for parallel growth | count | 1 |
| sphere_overlap | Sphere sampling factor along processes (centre spacing = $r/\text{sphere\_overlap}$) | – | 4 |
| hops_spheres_to_ignore | Neighbouring spheres along a process ignored in self-collision checks | count | 24 |
| *Skeleton* |  |  |  |
| skeleton_growth | Flag: grow from an input skeleton rather than *de novo* | bool | 0 |
| input_skeleton_file | Path to input skeleton file (if `skeleton_growth=1`) | – | (empty) |
| *Cell parameters* |  |  |  |
| soma_radius | Soma radius | $\mu\text{m}$ | 1 |
| vox_sizes | Cubic simulation-voxel side lengths ($x\ y\ z$) | $\mu\text{m}$ | 100 |
| nb_process_populations | Number of distinct process populations (separate parameter sets) | count | 1 |
| *Process: growth limits* |  |  |  |
| branching_order | Maximum branching (bifurcation) order of processes | count | 3 |
| *Process: number of primary processes* |  |  |  |
| nb_primary_processes_mean | Mean number of primary processes emanating from the soma | count | 6 |
| *Process: primary orientations* |  |  |  |
| oriented_processes | Flag: primary processes have a preferred orientation (vs. isotropic) | bool | 0 |
| process_orientations | Preferred orientation unit vector ($x\ y\ z$) | – | (0, 0, 0) |
| process_orientation_std | Angular spread about the preferred orientation | $^\circ$ | 0 |
| *Process: tapering* |  |  |  |
| tapering | Flag: enable radial tapering of processes | bool | 0 |
| tapering_in_secondary_processes | Flag: apply tapering to post-bifurcation processes | bool | 0 |
| tapering_radius_start | Radius at the start of tapering (mean) | $\mu\text{m}$ | 0 |
| tapering_radius_start_std | Std. of the start radius | $\mu\text{m}$ | 0 |
| tapering_length | Arc length over which tapering occurs (mean) | $\mu\text{m}$ | 0 |
| tapering_length_std | Std. of the tapering length | $\mu\text{m}$ | 0 |
| *Process: radius* |  |  |  |
| process_radius_mean_lognorm | Mean $\mu$ of the underlying normal of the log-normal radius distribution | $\log(\mu\text{m})$ | 0 |
| process_radius_std_lognorm | Std. $\sigma$ of the underlying normal (beading amplitude) | – | 0.1 |
| process_radius_correlation_length | Correlation length of radius fluctuations (beading) along a process | $\mu\text{m}$ | 5 |
| *Process: segment length* |  |  |  |
| segment_length_shape | Gamma-distribution shape $k$ of segment lengths | – | 1 |
| segment_length_scale | Gamma-distribution scale $\theta$ of segment lengths | $\mu\text{m}$ | 10 |
| segment_length_factor_B0 | Multiplicative segment-length reduction per branching order | – | 1 |
| *Process: branch angle* |  |  |  |
| branch_angle_mean | Mean angle between a child branch and its parent | $^\circ$ | 25 |
| branch_angle_std | Std. of the branch angle | $^\circ$ | 10 |
| *Process: undulation* |  |  |  |
| undulation | Flag: enable centreline undulation (wiggle) | bool | 0 |
| undulation_std | Amplitude (std.) of undulation displacement | $\mu\text{m}$ | 0.1 |
| undulation_correlation_length | Correlation length of undulation along a process | $\mu\text{m}$ | 5 |
| *Process: protrusions (spines)* |  |  |  |
| protrusions | Flag: add spine-like protrusions | bool | 0 |
| protrusion_head_radius_mean | Mean protrusion-head radius | $\mu\text{m}$ | 0.4 |
| protrusion_head_radius_std | Std. of the protrusion-head radius | $\mu\text{m}$ | 0 |
| protrusion_neck_length_mean | Mean protrusion-neck length | $\mu\text{m}$ | 1.25 |
| protrusion_neck_length_std | Std. of the protrusion-neck length | $\mu\text{m}$ | 0 |
| protrusion_neck_radius_mean | Mean protrusion-neck radius | $\mu\text{m}$ | 0.125 |
| protrusion_neck_radius_std | Std. of the protrusion-neck radius | $\mu\text{m}$ | 0 |
| protrusion_relative_volume_fraction | Target fraction of process volume in protrusions | – | 0.01 |

#### 2.1.1. Cell growth types

OCTOPUS comprises two types of cell growth that determine how the cell branching architecture is created: free growth and skeleton-based growth (Figure 1B). The free growth approach is designed as a minimal strategy, with morphological parameters sampled directly from input distributions without requiring a 3D cell reconstruction. This provides a simple framework that does not require additional inputs or data and allows a wide range of cellular morphologies to be generated from the same underlying distributions. However, because the growth process is not informed by a predefined morphology, this approach may fail to reproduce cells exhibiting atypical branching patterns. In contrast, skeleton-based growth uses a cell skeleton, which can be obtained from histological cell reconstructions, to shape the growth process and capture the overall branching architecture of the target cell. This reproduces highly specific cellular architectures, including complex branching patterns. However, this approach requires an existing cellular skeleton as an additional input. Moreover, because the generated cells inherit the branching architecture of the reference skeleton, the resulting morphological variability is inherently reduced. For both growth types, other cell geometry features of the grown cells, such as beading, undulation, protrusions, etc., are added in the same way, as described below.

In free growth, the cell is grown without a predefined structure, with random branch orientations and branching. To obtain a branch segment length compatible with the input configuration, new branches are preferentially grown at segment ends from which only one branch emerges. Free growth is stopped once it reaches the user-defined target branching order (number of consecutive branchings). During growth, the branching order of all processes is stored, starting at zero for the primary processes and incrementing after each branching. From the branching order of all grown processes, an average branching order is obtained, weighted by segment length. In the skeleton-based approach, the cell skeleton serves as a scaffold on which the cell generator adds morphological complexity. The skeleton can be either manually defined or derived from a 3D cell reconstruction using OCTOpool. The direction of the branches, their arborization, and their segment lengths are preserved, but other morphological features such as branch undulation and beading can be tuned.

#### 2.1.2. Soma

To start creating a cell, first the soma is placed. In free growth, it is positioned at the center of a cube of user-defined size. In skeleton growth, the skeleton defines its location within a cube defined by an edge length of twice the maximum path length from the soma derived from the skeleton, and adding a 10 % margin. The soma is approximated as a sphere, parametrized by its radius.

#### 2.1.3. Processes

Next, primary processes are grown from the cell soma. In free growth, the user specifies the number of primary branches, which emerge perpendicular to the soma’s surface. In skeleton growth, the number of primary dendrites is derived from the skeleton. As OCTOPUS represents cellular features by overlapping spheres, growing a process corresponds to generating subsequent spheres. The process radius is deterministically initialized to the mean of a log-normal distribution, to which tapering and beading can be added (see below). The sphere-to-sphere spacing is defined by an overlap factor, such that the center-to-center distance between consecutive spheres equals the radius of the preceding sphere divided by the overlap factor. Secondary branches are added iteratively. In free growth, when adding secondary branches, the branching angle between the secondary branch direction and the parent branch direction is sampled from a user-specified Gaussian distribution. In skeleton growth, the branching angle is obtained from the skeleton. Processes are grown in parallel, with each used CPU core growing one process. If one process has been successfully grown, overlap checks with all previous branches are performed, and only if no overlap is detected, is the branch added to the substrate. Once all processes are grown, the next growth iteration is initiated, and growth is stopped once the target branching order is reached.

##### Segment length

The growth of an individual process is stopped when a segment length is reached. In free growth, this length is drawn from a user-specified Gamma distribution, while in skeleton growth, it is obtained from the skeleton. Newly grown short processes overlap less often with preexisting processes than long processes. Hence, more short than long processes are added to the substrate, as they are less likely to be rejected due to overlap, introducing a potential bias in free growth process lengths. To avoid this bias, OCTOPUS tracks non-realized process lengths and grows processes with these lengths in the next process growth iterations. In some cells, the segment length strongly depends on the branching order, with more distal branches exhibiting progressively shorter lengths. To mimic this behavior, a parameter scales the drawn segment length according to the branching order of that segment. Specifically, the drawn segment length is multiplied by a factor *f*^BO^, where BO is the branching order.

##### Primary process orientation

In free growth, the user can specify a preferred orientation of primary processes to generate non-isotropic cells. A 3D vector specifies the preferred orientation, and a standard deviation controls how much the generated primary processes deviate from their preferred direction.

##### Tapering

Tapering describes the decrease of the process radius *r* dependent on the distance to the soma *x* (Figure 1A). The tapering is specified by the process radius at the soma (mean ± SD) and a characteristic length (mean ± SD). This behavior is described as an exponential decrease following *r*(*x*) = *r*_0_ + (*r_start_ - r*_0_) exp (−*x*/*l_tapering_*), where *r*_0_ is the mean process radius without tapering, *r_start_* the tapering start radius, and *l_tapering_* the tapering length. Another parameter determines whether tapering continues in secondary processes.

##### Ornstein-Uhlenbeck model of undulation and beading

Along a cell process, its direction and radius vary stochastically around a preferred direction and average radius, respectively. Prior research established that this variation is characterized by short-range disorder, implying independence of long-range variations (Novikov et al., 2014). We simulated these variations using the Ornstein-Uhlenbeck model (OU) as a minimal model for these characteristics (Uhlenbeck and Ornstein, 1930). The OU combines a random variation of a variable *ψ* with a parameter *x*, described by white noise *η*(*x*) and a linear relaxing force to a long-term mean µ.

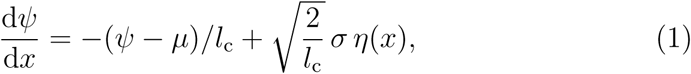

where *l*_c_ is the correlation length and *σ* the standard deviation of *ψ*.

##### Undulation

Undulation is implemented as an OU of the process tangent *t⃗* along the process length x (Figure 1A). As the tangent is a unit vector, the OU corresponds to 2D Brownian motion on the sphere with a relaxing force toward a preferred direction *t⃗*_0_. The preferred direction is drawn at the start of the process growth, respecting a user-specified branching angle distribution. We chose a relaxing force proportional to the angle between *t⃗* and *t⃗*_0_:

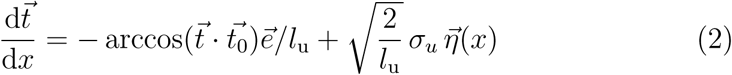

with *e⃗* the unit vector pointing along the geodesic toward *t⃗*_0_, and *η⃗*(*t*) being 2D white noise on the unit sphere. Thus, undulation is characterized by the direction correlation length *l_u_* beyond which the process direction becomes uncorrelated, and a parameter *σ_u_*, which for small angles corresponds to the standard deviation of the angle.

##### Beading

Beading of the process radius r along process length x was implemented as a lognormal OU model (Figure 1A). We chose this model, as the process radius is a nonnegative variable. Furthermore, our evaluation of the replicated histological cell reconstructions showed that a lognormal distribution described the radius distribution slightly better than a gamma distribution (not shown). Thus, the evolution of the logarithm of the radius r_log_ was modeled by

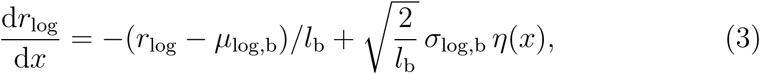

where *µ*_log,b_ and *σ*_log,b_ are the mean and standard deviation of *r*_log_ and *l*_b_ the beading correlation length. To disable beading, *σ*_log,b_ can be set to zero.

##### Process populations

Some cells have qualitatively different types of processes. For instance, pyramidal neurons have apical and basal dendrites, which differ not only in their growth direction but also in their process length and tapering. In free growth, OCTOPUS can grow cells with a user-defined number of process populations. For each process population, all process parameters must be defined separately. The process populations are grown sequentially, obeying the order of the user-specified parameters. This option is not currently supported in the skeleton-based approach.

#### 2.1.4. Protrusions

Protrusions on cell processes (Figure 1A), such as dendritic spines and astroglial leaflets, are approximated by a cylindrical neck that starts on a branch and a spherical head placed on top of the neck, similar to (Ofer et al., 2021; Şimşek et al., 2025; Chakwizira et al., 2025). For growing a protrusion, its start coordinate on the cell process and the growth direction are randomly sampled. The protrusions are grown until they reach the userdefined protrusion volume fraction relative to the branch volume.

#### 2.1.5. Implementation of the collision check

Collision checks are performed once a branch has finished growing. To speed up the collision checks between growing branches and existing elements, two look-up tables (LUTs) are defined. First, a 3D LUT encodes the validated elements’ positions in a bitwise manner. The resolution of the LUT is determined by its size, a tunable parameter limited by the computer memory available. We chose a size of 1000 1000 (1000/8), which for a cubic voxel of 100 µm, leads to a resolution of 0.1 µm. Second, another 3D LUT called “sphere map” stores the validated elements’ indices. Collisions between a growing branch and existing elements are first checked with the bit-wise LUT. This check consists of simply accessing the LUT entries overlapping with the elements to check if they are occupied. If the LUT is free, the element can be safely added to the growing substrate. If not, further collision checks are necessary. Potentially colliding spheres are retrieved from the sphere map, and overlaps are checked against all spheres in the grown branch. If no overlap is detected, the branch is added; otherwise, it is discarded.

In some cases, overlap between branches may be needed for an accurate representation of the cell. For example, in areas of process branching, the initial spheres of the two branches emerging from the same origin will overlap. In beaded and tapered processes, more consecutive spheres may overlap than specified in the configuration file. To exclude these collisions from the overlap check, a parameter defines how many connected spheres should be ignored in the overlap check. The hops are counted across branchings and the soma as well.

### 2.2. OCTOpool: cell analysis, generation and Monte Carlo simulation

For convenient use of OCTOPUS in modern data analysis pipelines, we developed the OCTOpool toolbox in Python. OCTOpool provides submodules to generate, analyze, and visualize the cells, run Monte Carlo (MC) simulations of diffusion within the generated cells, and analyze the walker trajectories and the simulated dMRI signals from MC simulations. In the Supplementary Methods, we describe the functionality central to the presented results, while extensive documentation is included in the OCTOpool repository, which will be released upon publication.

### 2.3. Virtual experiments

We performed several virtual experiments to validate the biological plausibility of OCTOPUS-generated cell geometry features and to demonstrate the capabilities of our toolbox. Water diffusion within digital cell substrates was simulated using a customized version of the MCDC simulator that is compatible with substrates represented by overlapping spheres (Rafael-Patino et al., 2020; Nguyen-Duc et al., 2026a). In experiment 1, we tested the compatibility of OCTOPUS-generated undulation and beading with the short-range disorder property that was previously established for fine-grained cellular heterogeneities. In experiment 2, we further underscored the plausibility of OCTOPUS cell growth by reproducing the 1/√*b* power law of the dMRI signal at high b values and analyzing the impact of the cell soma on this relationship. In experiment 3, we demonstrated the versatility of OCTOPUS by reproducing eight different cell types, including neurons and glial cells, and analyzed the accuracy and limitations of the implemented cell growth algorithms. In experiments 4 and 5, we showed that OCTOPUS enables the study of the effects of process branching and dendritic spines, reproducing previously reported behavior.

If not otherwise stated, the free diffusivity was set to 2 µm^2^/ms, the random-walk step size to 0.2 µm. We compared the resulting dMRI signals quantitatively by calculating their maximum signal deviation max*_b_*∥*S*_1_ – *S*_2_∥ across all simulated b values, and reported values in percent units with respect to the signal at b = 0 for better readability. Computations were carried out on a high-performance computing cluster using a single node (2 × 24 cores, 1.5 GHz).

#### Experiment 1: Time-dependent diffusion in elongated processes

To assess the biological plausibility of the beading and undulation models of OCTOPUS, we studied the time dependence of the diffusivity D and kurtosis K in substrates consisting of a single cell process. Beading and undulation are expected to introduce short-range disorder, wherein D and K should follow a functional form of 1/√*t* at long diffusion times (Novikov et al., 2014). Simulations were conducted on three substrates, each comprising a 1.5 mm long process with a radius of 1 µm, with straight, undulated, and beaded morphologies, respectively. The undulated process had an undulation SD of 0.3 and a correlation length of 10 *µ*m. The beaded process was parametrized with *σ*_log,b_ of 0.3 µm and a correlation length of 5 µm. The MC simulations were run for a maximum time of 500 ms and 10^6^ walkers, leading to a walker density of 212 µm*^−^*^3^. Walkers were initialized in a central region within the voxel, leaving a 220 µm buffer at both extremities to avoid edge effects. The parameters D and K were calculated from the walker’s trajectory using the OCTOpool toolbox, following equations 2 and 3 in (Fieremans and Lee, 2018). For the straight neurite, the mean diffusivity and kurtosis were reported. For the undulated and beaded neurites, the functional form of short-range disorder (*S* = *a* + *bt*^−0.5^) was fitted (Novikov et al., 2014), and the coefficient of determination or variance explained r^2^ was reported.

#### Experiment 2: High b stick power law in the presence of a soma

We tested whether the powder-averaged dMRI signal from generated cells follows the 1/√*b* power-law that was found experimentally in white matter (Veraart et al., 2019). Similarly to Olesen et al. (2022), we evaluated the impact of the soma on the intracellular diffusion signal in a neuron, as well as the influence of cellular geometry. We performed MC simulations in six distinct substrates: three comprising full cells (soma and processes) and three comprising processes only. Each substrate consisted of six processes, each 70 µm long with a radius of 0.69 µm. In the cell substrates, the soma had a radius of 4 µm (corresponding to astrocyte somata (Aird-Rossiter et al., 2026a)). For all substrates, straight, undulating, and beaded process morphologies were evaluated. The undulation had an SD of 0.2 and a correlation length of 5.7 µm, while the beading was parametrized with σ_log,b_ of 0.3 and a correlation length of 5 µm. For each substrate, dMRI signals were computed in five independently-grown replicas using a pulse-gradient spin echo sequence, with *δ* = 4.5 ms, Δ = 16 ms, echo time of 21 ms (equivalent to simulation time), *b* = 0 to 10 ms/µm^2^ in steps of 1 ms/µm^2^, and *b* = 10 ms/µm^2^ to 100 ms/µm^2^ in steps of 10 ms/µm^2^, and 128 uniformly distributed gradient directions, mimicking the parametrization used in (Olesen et al., 2022). Simulations were run using 10^5^ walkers, leading to walker densities of more than 112 µm*^−^*^3^. To estimate the power law exponent, we performed a linear fit in log-log space.

#### Experiment 3: Histological cell reproduction

To analyze the versatility and limitations of the OCTOPUS toolbox, we reproduced histological cell reconstructions using the OCTOpool cell reproduction feature.

To this end, we obtained eight openly available cell meshes from Aird-Rossiter et al. (2026a) for eight cell types: astrocytes, basket cells, GABAergic neurons, glutamatergic neurons, granule cells, microglia, Purkinje cells, and pyramidal neurons. In most cases, we picked the first cell in the repository, preferentially using rat cells. In some cases, we used later cells in case they had higher morphological complexity and higher reconstruction resolution, indicated by finer features in the cell reconstruction (microglia, for instance).

We analyzed the properties of these original histological reconstructions using OCTOpool and recreated similar cells using the free growth and skeleton growth modes of OCTOPUS. For each original cell and growth mode, we created 10 cell replicas to assess their statistical variability. Cell geometry was quantitatively analyzed using the OCTOpool toolbox and by obtaining Sholl analyses for each cell.

MC simulations were conducted in the histologically segmented cells given as .swc files, which we converted to a format of overlapping spheres mimicking the OCTOPUS substrate format, as well as their corresponding freely and skeleton-based grown counterparts. We used similar b-values to the ones used in experiment 2, but tested Δ = 25.5 ms to 105.5 ms, in steps of 10 ms, with *δ* = 16.5 ms and a simulation time of 127 ms. The simulations were run using 10^5^ walkers, ensuring a walker density > 1 µm*^−^*^3^. We validated that the predicted dMRI signals have a maximum average deviation of 2.6 % from simulations with 10^6^ walkers, while the mean average deviation was 0.4 % for all eight cell types and Δ.

#### Experiment 4: Signature of branching

To quantify the effect of branching, we regenerated cells using a modified version of the glutamatergic neuron reproduction in Experiment 3. First, to increase the signal’s sensitivity to the cell processes, we removed the tapering of the glutamatergic neuron. Moreover, we effectively removed the soma by reducing its size to the average process radius. Next, we used the resulting configuration to freely generate a branched glutamatergic neuron. To create a non-branched glutamatergic neuron with the same volume as the branches, we calculated the total combined length of the processes of the branched neuron and divided it by the number of primary processes. Then we generated a cell with only primary processes, each of length as calculated, resulting in the same process volume, as the average process radius was the same.

We performed MC simulations in the branched and non-branched substrates to assess the impact of diffusion at long times as proposed before (Palombo et al., 2016), wherein metabolite diffusion was studied up to a time of 2 s. As metabolite diffusivity is approximately four times lower than our chosen free diffusivity (2 µm^2^/ms), we simulated diffusion at Δ = 500 ms (corresponding to 2 s diffusion time for metabolites). We simulated *b* = 1 ms/µm^2^ to 10 ms/µm^2^ in steps of 1 ms/µm^2^.

#### Experiment 5: Signature of protrusions

Recently, the effect of spines on the dMRI signal has been studied (Şimşek et al., 2025; Chakwizira et al., 2025). We examined this effect by using the reproduction of the glutamatergic neuron from Experiment 3, to which we added spines following the parametrization proposed in Şimşek et al. (2025); Chakwizira et al. (2025). The spines were defined by a protrusion head radius of 0.4 µm, a neck length of 1.25 µm, and a neck radius of 0.125 µm. The protrusion volume fraction relative to the branch volume was set to 40 %.

For the MC simulation, we decreased the step length to 0.02 µm to be sufficiently smaller than the smallest spine dimension. We simulated b = 1 ms/µm^2^ to 10 ms/µm^2^ in steps of 1 ms/µm^2^.

## 3. Results

### 3.1. OCTOPUS and OCTOpool generate and replicate cells in seconds

On a desktop computer (3.2 GHz, 32 cores), almost all substrates used in this work were generated in less than 15 s. The only exception was the generation of the glutamatergic neuron with spines, which took 3 min.

OCTOpool cell replication, including a complete analysis of a given 3D cell reconstruction, took on average (13 ± 30) s for ten replicas grown either freely or skeleton-based. Notably, cell replication took less than 15 s across all configurations except the freely grown pyramidal neuron, which took 2 min.

### 3.2. Beading and undulation induce biologically plausible time-dependent diffusion

To check the biological plausibility of OCTOPUS substrates, we investigated the time dependence of *D* and *K* within straight, undulated, and beaded processes (Fig. 2). As expected, *D* and *K* in the straight branch showed no time dependence. The observed *D* corresponded to the input free diffusivity of 2 µm^2^/ms, and K was close to zero. In contrast, the undulated and beaded processes showed signatures of short-range disorder, with *D* and *K* scaling as 1/√*t*. The diffusivity diminished over time, with an excellent power law fit for *t* > 100 ms. Kurtosis initially increased with diffusion time, before decreasing again with power law scaling.

**Figure 2:**
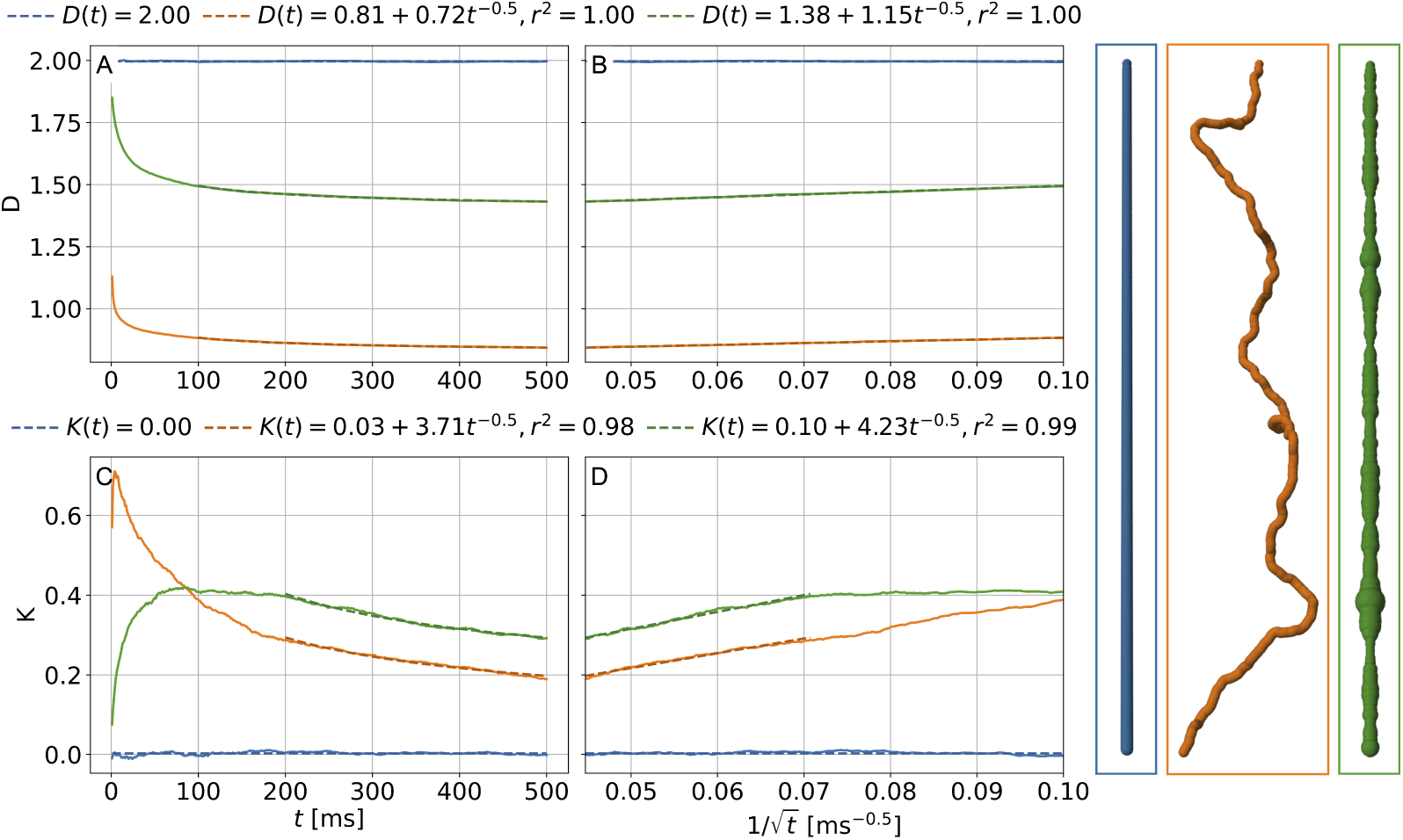
Diffusivity (*D*) and kurtosis (*K*) time dependence in the intracellular space of single processes. Processes with three geometries are shown: straight (blue), undulated (orange), and beaded (green), each of length 1.5 mm. Power-law curves *y* = *a* +*bt^−^*^0.5^ were fitted to *D* and *K* of beaded and undulated processes, and the coefficients of determination, or variance explained, *r*^2^ are shown. The dashed lines show the range of diffusion times used for fitting.

### 3.3. Substrates show stick power law at high b-values

Furthermore, we examined the behavior of the dMRI signal at high *b*-values, which has been reported to scale with 1/√*b* (Veraart et al., 2019; Olesen et al., 2022). We generated cells with straight, undulated, and beaded branches, and also regenerated the same substrates without a soma (Fig. 3). As expected, the intracellular signal from branches-only substrates agreed with the stick power law for almost the entire range of b-values. The fitted exponents were 0.51, 0.52, and 0.51 for the straight, undulated, and beaded cases, respectively, hence agreeing with the stick power law exponent of 0.5 with an accuracy as reported previously (Olesen et al., 2022). When adding a soma, the diffusion signals deviated from the power law for *b* < 30 ms/µm^2^, while approximating the power law for higher *b*, where the soma contribution is effectively suppressed. Hence, dMRI signals obtained from diffusion within OCTOPUS-created substrates show plausible high-b behavior.

**Figure 3:**
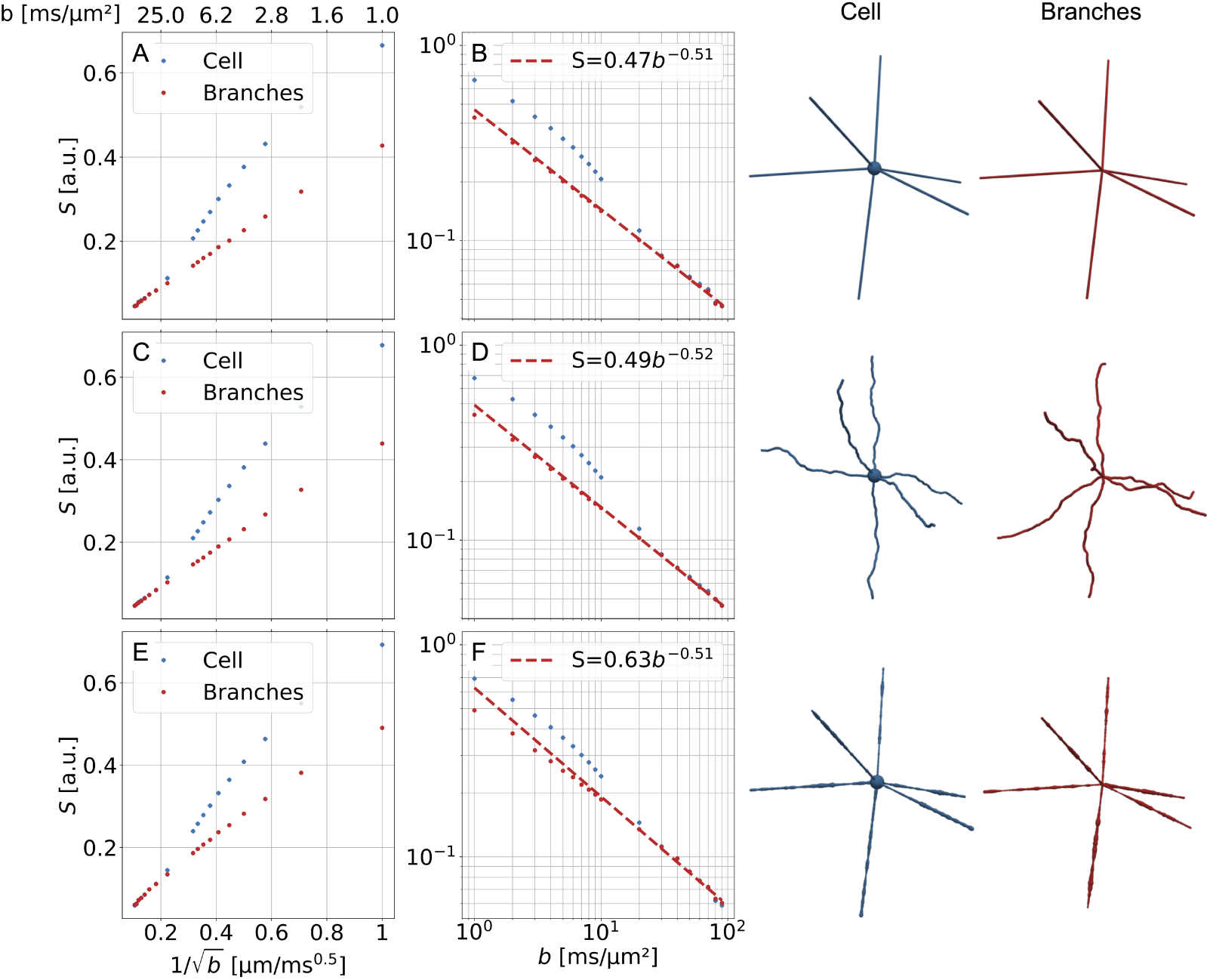
Stick power law for high *b*-values in generated cells. dMRI signals are shown for cells including both branches and cell bodies (blue), as well as for branches only (red). For both cases, three geometrical morphologies were obtained: straight, undulated, and beaded. Linear fits in log–log space are overlaid. Example cells used for the simulations are shown in the right columns.

### 3.4. OCTOpool replicates histological cells

To showcase the flexibility of OCTOPUS and OCTOpool, we duplicated histological cell reconstructions of eight different types.

We generated these cells using both free and skeleton-based growth (Figs. 4, 5). With the skeleton-based growth approach, cells preserve the original branching architecture, segment lengths, and branching angles. Additional input parameters for geometrical features, such as beading and undulation, were estimated from the original cells. As a result, these generated cells strongly resemble the original morphologies, while allowing less variability than in free growth. In contrast, the free growth approach does not rely on any predefined architecture. Instead, cells are generated solely by sampling parameters from input distributions. The same input parameters for geometrical features as for skeleton-based growth were used. In Supplementary Tables S1 and S2, we reported all OCTOpool-estimated cell geometry parameters. As an illustrative example, we reproduced the values for the glutamatergic neuron in Table 2. Most of the parameters agree well between original cells and OCTOPUS recreations, underscoring the ability of the estimators implemented in OCTOpool to assess cellular geometry. Nevertheless, beading and undulation correlation lengths were often underestimated in generated cells compared to original ones, as the correlation lengths were often comparable to or greater than the segment lengths (Table 2). Moreover, estimations of beading and tapering impact each other, as both describe processes affecting the branch radius, rendering a faithful tapering quantification in the presence of pronounced beading difficult.

**Figure 4:**
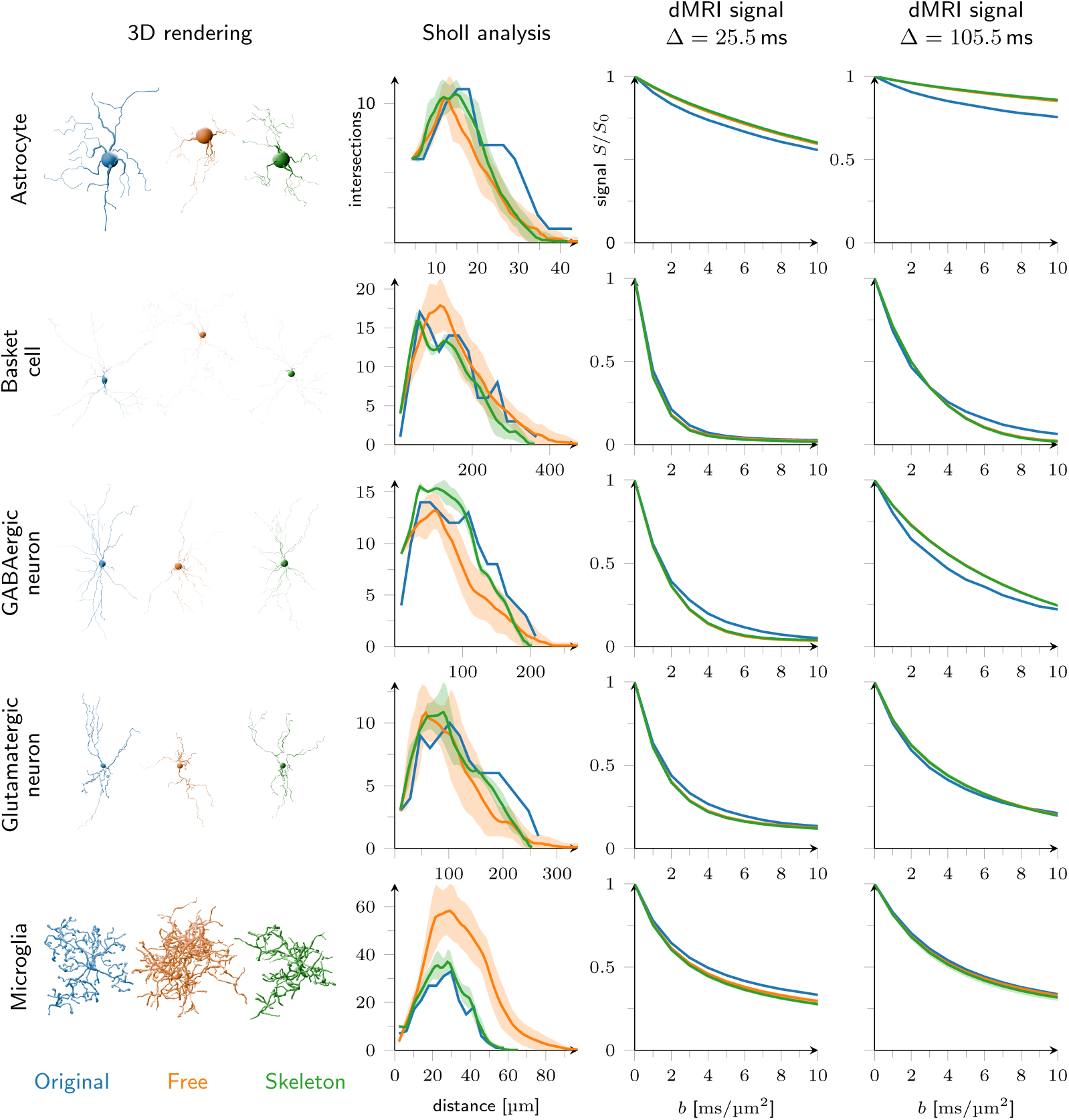
Cell reproductions where free and skeleton-based growth agree. Each row shows a different cell type. From left to right: histological reconstruction (original, blue), free growth (free, orange), skeleton-based growth (skeleton, green), the corresponding Sholl analyses, and dMRI signal decays for short (25.5 ms) and long 105.5 ms diffusion times. For Sholl analyses and dMRI signal plots, the mean across ten grown cells is shown with the standard deviation as a shaded area. For all displayed cell types, freely and skeleton-based grown cells show only minor deviations in their Sholl analysis and dMRI signal decays, indicating that free growth captures the geometry of these cells accurately.

**Table 2:** Reproduction of a glutamatergic neuron. Values in the Free and Skel. columns are mean *±* standard deviation over 10 repetitions.

| Parameter | Orig. | Free | Skel. |
| --- | --- | --- | --- |
| Number primary dendrites | 3 | 3 | 3 |
| Soma radius ($\mu\text{m}$) | 10.57 | 10.57 | 10.57 |
| Branching order | 2.91 | $2.83 \pm 0.06$ | 2.50 |
| Branching angle mean ($^\circ$) | 46.0 | $56.5 \pm 2.2$ | $59.5 \pm 5.6$ |
| Branching angle std ($^\circ$) | 31.5 | $29.3 \pm 2.5$ | $33.2 \pm 3.7$ |
| Segment length mean ($\mu\text{m}$) | 51.63 | $45.67 \pm 7.80$ | $56.12 \pm 0.02$ |
| Branch radius mean ($\mu\text{m}$) | 0.35 | $0.43 \pm 0.06$ | $0.46 \pm 0.05$ |
| Branch radius std. ($\mu\text{m}$) | 0.11 | $0.09 \pm 0.02$ | $0.10 \pm 0.01$ |
| Beading $\mu_{\log}$ | -1.10 | $-0.87 \pm 0.16$ | $-0.80 \pm 0.12$ |
| Beading $\sigma_{\log}$ | 0.30 | $0.20 \pm 0.02$ | $0.22 \pm 0.02$ |
| Beading $l_c$ ($\mu\text{m}$) | 28.95 | $8.57 \pm 1.93$ | $11.31 \pm 1.47$ |
| Tapering $r_0$ ($\mu\text{m}$) | 1.59 | $1.76 \pm 0.20$ | $1.74 \pm 0.32$ |
| Tapering length ($\mu\text{m}$) | 122.72 | $76.43 \pm 21.78$ | $70.94 \pm 25.42$ |
| Undulation $\sigma$ | 0.26 | 0.26 | 0.26 |
| Undulation $l_c$ ($\mu\text{m}$) | 16.61 | $9.60 \pm 1.45$ | $10.78 \pm 1.09$ |

For most cell types, the free growth approach is sufficient to reproduce histologically accurate morphologies. Among them are astrocytes, basket cells, GABAergic and glutamatergic neurons, and microglia (Fig. 4). For all of these cells except for microglia, free and skeleton-based Sholl analyses agreed overall well, although not always within the standard deviation across cell generation repetitions. For microglia, the 3D rendering and the Sholl analysis both indicate less dense processes in the original and skeleton-based cells. The maximum dMRI signal deviation between freely and skeleton-based grown cells was (0.8 ± 0.5) %, indicating good agreement between the two approaches at the level of the generated dMRI signal. We observed some residual deviation between the dMRI signals of original and reproduced cells (maximum signal difference of (5 ± 3) %), indicating imperfections in cell reproduction. Nevertheless, the simulated dMRI signal between different cell types showed more pronounced differences (maximum signal difference of (33 ± 19) %) than the error between the original and reproduced cells. This implies that the dMRI signal still carries a signature that distinguishes these cell types.

For others, particularly anisotropic cells, free growth failed to reproduce the cellular architecture. We observed this in granule cells, Purkinje cells, and pyramidal neurons (Fig. 5). These deviations between original and freely grown cells were apparent both in the Sholl analyses and dMRI signal decays (maximum signal difference between freely and skeleton-based cells was (4.6±1.0) % for the granule cell, (18±3) % for the pyramidal neuron, across the diffusion times). For the granule cell and the pyramidal neuron, the skeleton-based substrates alleviated these differences, and dMRI signals corresponded much better to the original cells (maximum signal difference of (1.1 ± 0.2) % for the granule cell, (3.5 ± 0.3) % for the pyramidal neuron). For the Purkinje cell, OCTOpool overestimated undulation, leading to a decreased cell span in both free and skeleton-based growth, which is apparent by strongly differing Sholl analyses. Nevertheless, the dMRI signals agreed reasonably between all three cell reconstructions ((6.1 ± 1.7) % maximum signal difference across all reconstructions and diffusion times). To achieve better geometrical correspondence for all these cells, further manual tuning of OCTOPUS parameters is needed to achieve an accurate representation. This can be achieved by declaring several process populations and assigning growth parameters to each population individually. An illustration of manual tuning on a pyramidal cell is depicted in Fig. 6, where the apical and basal dendrites were defined as two distinct process populations.

**Figure 5:**
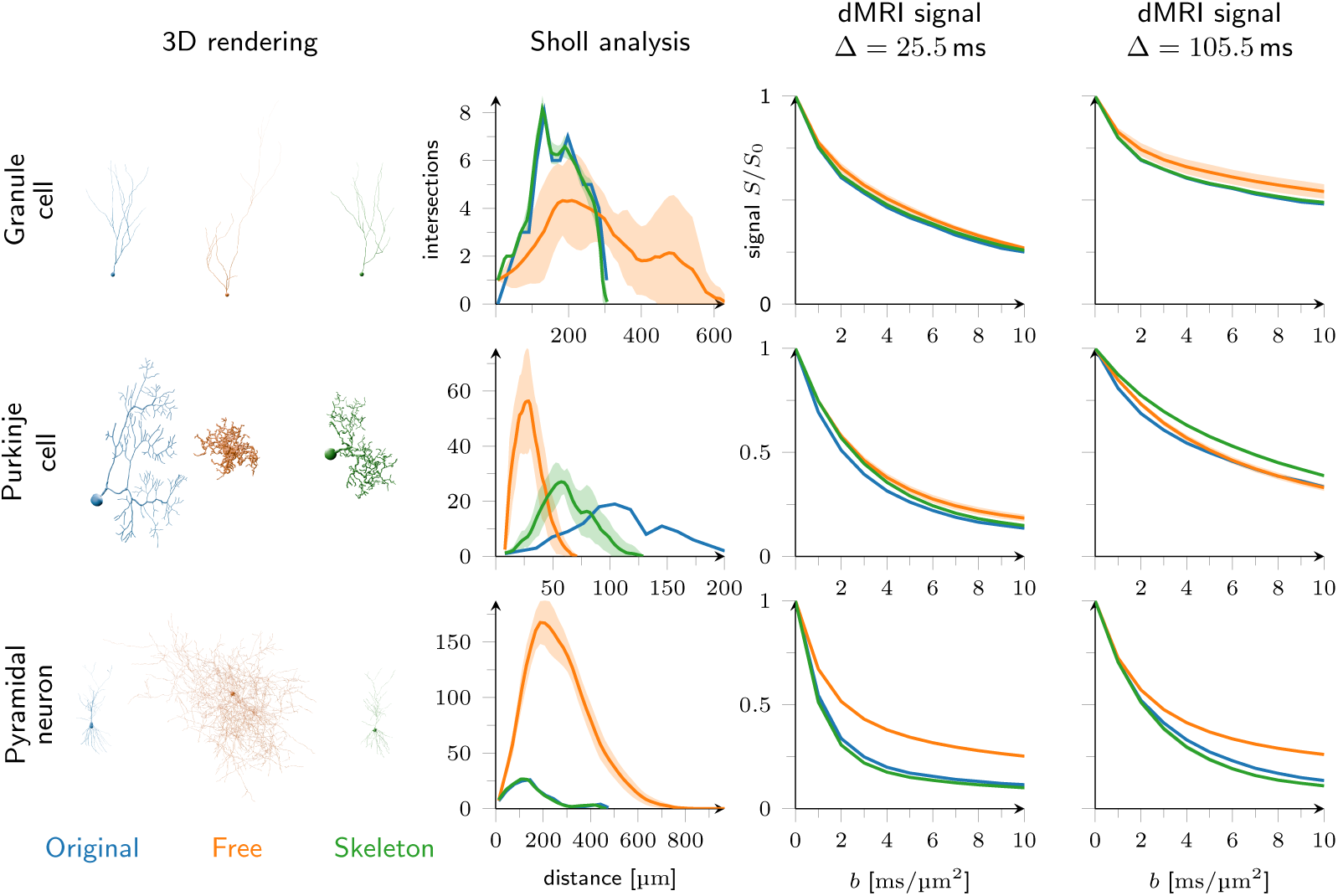
Cell reproductions where free and skeleton-based growth differ. Each row shows a different cell type. From left to right: histological reconstruction (original, blue), free growth (free, orange), skeleton-based growth (skeleton, green), Sholl analyses, and dMRI signal decays for short (25.5 ms) and long 105.5 ms diffusion times. All displayed cells show marked differences in their Sholl analyses between the original and grown cells. The dMRI signal reflects these deviations to a varying degree.

**Figure 6:**
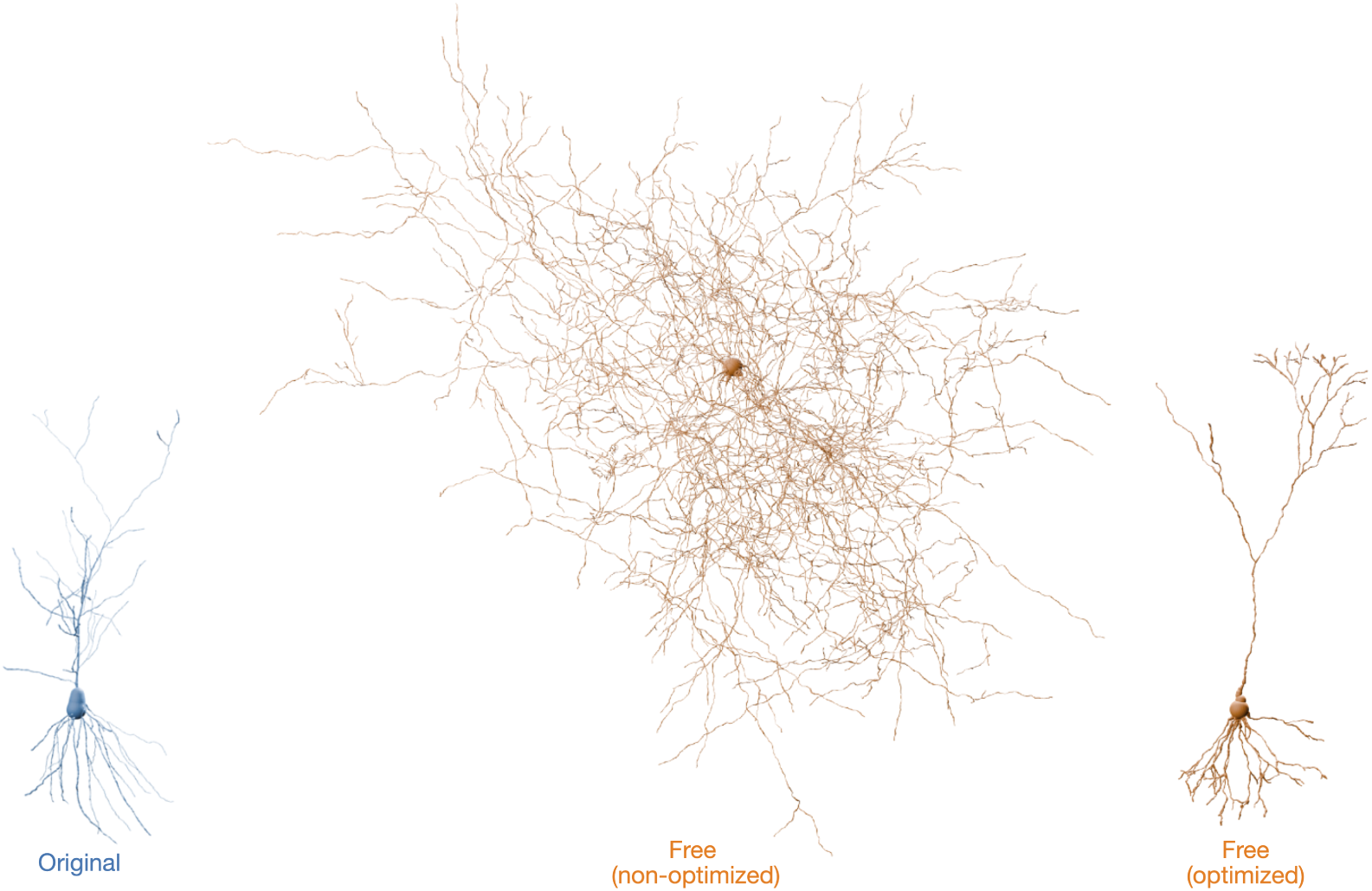
Illustration of manual optimization of the free-growth parameters of a pyramidal neuron. The original cell and the free (non-optimized) cell are taken from the pyramidal neuron in Figure 5. The free (optimized) cell uses two process populations, one for the apical and one for the basal dendrites. The process orientations are set to [0, 0, 1] and [0, 0, -1], and the process orientation standard deviations are set to 0.1 and 0.5, respectively.

### 3.5. dMRI signal shows signatures of branching at long diffusion times

Past research suggested that the metabolite diffusivity quantified by dMRS is sensitive to the cell branch length (Palombo et al., 2016). We reproduced this effect by generating freely grown glutamatergic neurons with and without branching, while reducing the soma size and removing tapering to increase sensitivity to branching effects (Fig. 7). The simulated dMRI signals showed the previously reported effect of branching: For a short diffusion time (10 ms, corresponding to a one-dimensional diffusion length of approximately 6 µm), the signals from both substrates show a minute maximum signal difference of 0.8 %. Here, the diffusion length is much shorter than the segment length of the branched substrate (l ≈ 45 µm), thus limiting the effect of branching. For a long diffusion time of 500 ms (corresponding to a diffusion length of 45 µm ≈ l), the decay of the signal from the branched substrate is slower, leading to an increased maximum signal difference of 3.8 %, indicating an effectively decreased diffusivity due to branching.

**Figure 7:**
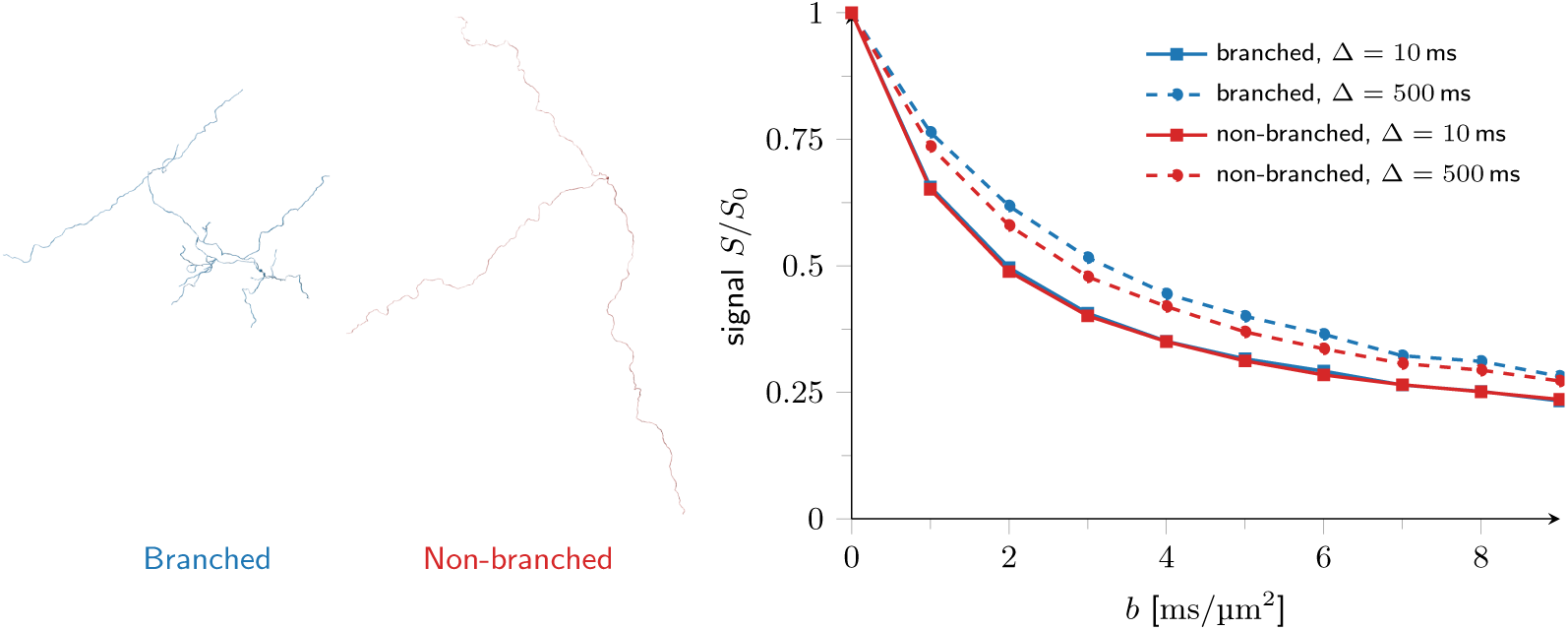
The effect of cell process branching on the dMRI time dependence. On the left, branched and non-branched freely grown glutamatergic neurons are shown. For the nonbranched substrate (red), the branch length was extended to result in the same total branch length as for the branched substrate (blue). On the right, the dMRI/S signal originating from these cells is shown at different diffusion times. At a short diffusion time (10 ms, filled lines), the signals from both substrates coincide, while at a long diffusion time (500 ms, dashed lines), branching reduces the effective diffusivity, as previously reported (Palombo et al., 2016).

### 3.6. dMRI signal shows signatures of protrusions

Recent research examined the potential effect of protrusions, in particular, dendritic spines, on the dMRI signal (Şimşek et al., 2025; Chakwizira et al., 2025). We used OCTOPUS to investigate this effect by generating glutamatergic neurons (parameters according to Table 2) with and without protrusions (Fig. 8). Between the cell with spines and the average signal from freely generated glutamatergic neurons, the dMRI signal showed a maximum difference of 37 %, much larger than the standard deviation between signals from freely grown glutamatergic neurons, which was (0.5±0.2) %, indicating a pronounced effect of the added spines. The signal from the spiny neuron was higher than the one from the non-spiny neuron, indicating that the spines decreased effective diffusivity, as water can be trapped in spines, which stops diffusion along the length of the dendrite. These results indicate that OCTOPUS is suitable for studying the effect of protrusions in realistic cells.

**Figure 8:**
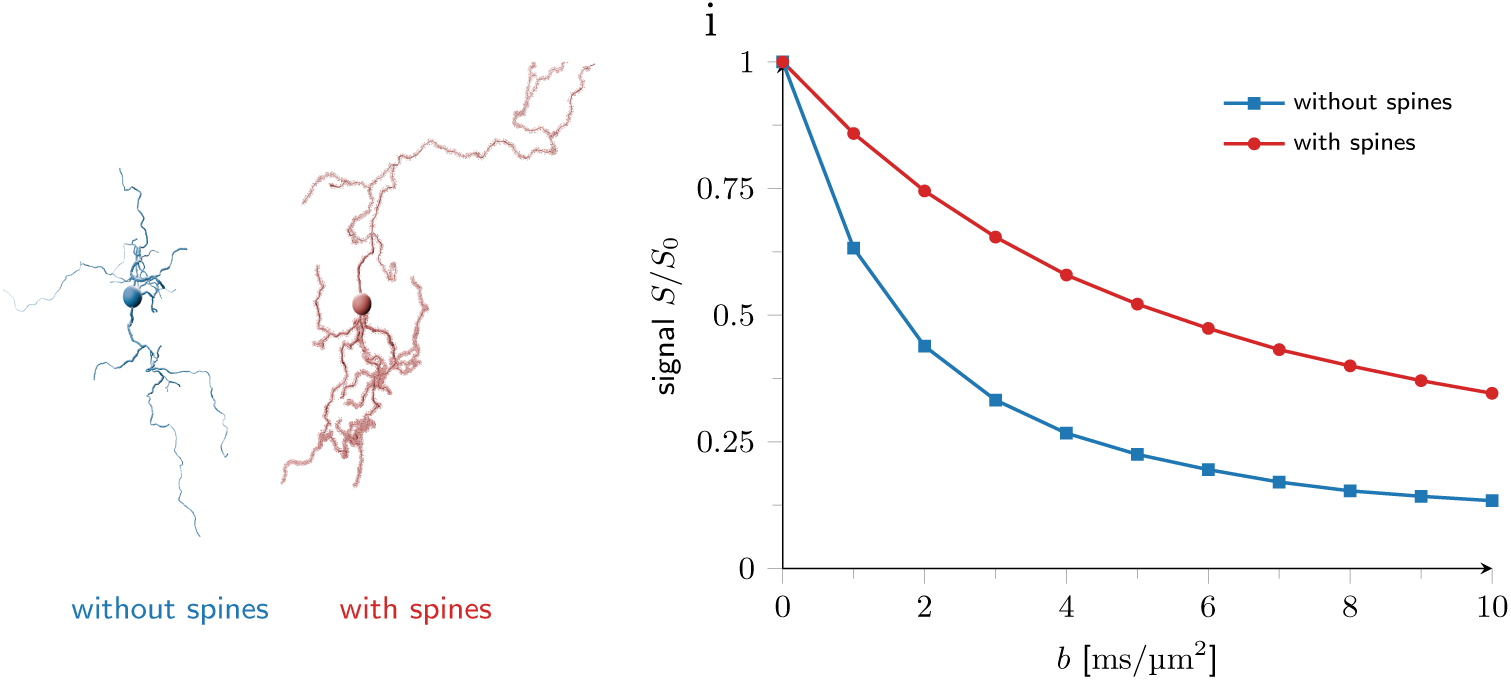
To assess the impact of protrusions on the dMRS/I signal, we freely generated glutamatergic neurons with and without spines (left). A markedly less pronounced signal decay was observed for the spiny glutamatergic neuron (right), indicating that spines hinder diffusion within the cell, as has been proposed in recent research (Şimşek et al., 2025; Chakwizira et al., 2025).

## 4. Discussion

This paper introduces OCTOPUS, an open-source C++ software capable of quickly generating a wide variety of brain cells with tunable morphological complexity (Fig. 1). We demonstrated the biological plausibility of the generated cells at multiple levels. First, for irregularities along elongated cell processes (due to undulation and beading), we reproduced established properties of dMRI signal: diffusion-time dependence compatible with short-range disorder (Fig. 2) and the stick power law at high b-values (Fig. 3). Second, we showcased OCTOPUS’s ability to reproduce a wide range of cell types with high agreement between input and output cell geometry parameters, such as soma and process radii, branching order and length, tapering, undulation, and beading. We compared their diffusion signals and Sholl analyses with histological cells (Figs. 4, 5), and analyzed how different cell growth strategies, free and skeleton-based growth, can mimic different cell geometries. For more isotropic cells, both skeleton-based and free-growth approaches generated similar signal decays. For more anisotropic cells such as pyramidal cells and granule cells, the skeleton prior was needed for a more faithful reconstruction. We showed how free growth can be adapted to accommodate more complex cell geometry (Fig. 6). Eventually, we demonstrated signatures of branching (Fig. 7) and spines (Fig. 8) in OCTOPUS-generated cells. By efficiently growing plausible cells, OCTOPUS contributes to the broader effort of building realistic brain tissue substrates.

### OCTOPUS generates biologically plausible cell processes

We first corroborated the biological plausibility of the OCTOPUS irregular process microstructure by finding a time dependence consistent with short-range disorder. In white matter axons, D and K are known to exhibit the signature of short-range disorder, a 1/√*t* time dependence (Novikov et al., 2014; Nguyen-Duc et al., 2026a). This confirmed that no artificial periodicity was introduced with the Ornstein-Uhlenbeck processes describing undulation and beading, which would alter diffusion and kurtosis time dependence (Novikov et al., 2014; Lee e t al., 2020a). Moreover, OCTOPUS-generated cell processes showed the 1/√*b* power law (Veraart et al., 2019), which was then altered by the addition of a soma. We obtained the stick power law for beaded and undulated processes, similar to previously published simulations in branches represented as perfect cylinders (Olesen et al., 2022).

Hence, we were able to show that OCTOPUS generates plausible complexity and irregularity of cellular processes.

To replicate cell geometries, OCTOPUS supports two growth types: free and skeleton-based. Interestingly, these growth types produced similar cells for some more isotropic cell types (Fig. 4), with Sholl analyses and dMRI signal agreeing well, while for other, more anisotropic cell types (Fig. 5), marked differences were observed. This indicates that more isotropic cells, including astrocytes, glutamatergic neurons, and GABAergic neurons, grow generically: their processes branch at intervals from the cell body, and the branch lengths follow the same probability distribution regardless of soma distance. Other cells (microglia, granule and Purkinje cells, pyramidal neurons) exhibit more complex and cell-specific patterns of process growth, leading their free and skeleton-based growth replicas to disagree. The increased maximum intersections in the Sholl analysis of freely grown microglia suggest that the original microglial processes do not branch as often, indicating that branching occurs preferentially in some processes. Furthermore, the increased maximum cell extent of freely grown microglia, granule cells, and pyramidal neurons may indicate that segment lengths between more branched and less branched processes are not the same. While different growth strategies are warranted to replicate cells of different types, the differences between free and skeleton-based replicas could be used to inform research on the functional relevance of the deviating growth patterns.

The free growth of pyramidal neurons was further refined to better resemble histological cells by introducing two distinct cell populations, corresponding to the apical and basal trees. However, the morphological parameters extraction step of OCTOpool does not support extracting parameters for multiple process populations. A more natural implementation of a framework capable of producing anisotropic cell morphologies will therefore constitute an important direction for future work, as presented in Palombo et al. (2019); Aird-Rossiter et al. (2026a).

### A novel tool for numerical studies of microstructure imaging

Previous studies have devoted considerable attention to developing white matter substrates composed of densely packed axons (Ginsburger et al., 2019; Callaghan et al., 2020; Villarreal-Haro et al., 2023; Nguyen-Duc et al., 2026a; Winther et al., 2024). In gray matter, however, the high morphological complexity of cells substantially increases the challenge of: i) generating individual cells that agree with histology, and ii) packing these cells within a voxel while preserving biologically plausible morphologies. Previous work has explored generating realistic gray matter cells, although the associated framework was not released as open-source (Palombo et al., 2019), or focused exclusively on astrocytes (Ginsburger et al., 2019; Nguyen-Duc et al., 2026a). More recently, Aird-Rossiter et al. (2026b) presented a framework for generating cellular substrates that capture the overall architecture of realistic cortical columns. However, their implementation does not allow explicit control over undulation, beading, or tapering, nor does it include protrusions. Hence, OCTOPUS fills a gap in the current landscape of synthetic tissue generators by providing an open-source, versatile framework generating multiple classes of realistic gray matter cells, with tunable and reproducible branching, beading, undulation, tapering, and protrusions.

Our results show how OCTOPUS enables studying the impact of complex cell geometry, such as beading, undulation, and protrusions, on dMRI signals. As anticipated, increasing geometrical complexity reduced diffusivity due to increased geometric hindrance (Ianus et al., 2021). At longer diffusion times, water molecules explore their environment further and reflect from cell boundaries, decreasing D. In parallel, irregularities initially elevate the kurtosis by introducing greater heterogeneity in diffusivity, but ultimately decrease K once substantial mixing of water or molecules occurs, as previously described (Lee et al., 2025). By controlling features such as beading and undulation, branching, protrusions, or soma size, OCTOPUS allows examining how these geometrical properties impact estimates of analytical biophysical models with simplified geometries. Down the road, OCTOPUS can also assist in establishing diffusion encoding schemes that sensitize the signals to specific microstructural features and in developing numerical biophysical models trained on OCTOPUS synthetic datasets.

OCTOPUS may play a particular role in dMRS, where simulations of intracellular metabolite diffusion, including increased cellular complexity, may enable the extraction of such features from experimental signal decays. Unlike dMRI, dMRS is primarily sensitive to the intracellular space, hence avoiding the step of packing cells into a dense substrate.

Another potential application of OCTOPUS could be to recreate histological cell reconstructions of healthy tissue, while selectively modifying geometrical features known to be altered in disease. For example, OCTOPUS would allow for studying the impact of reduced spine density on the dMRS signal, which was found in Purkinje cells in a rat model of creatine-transporter deficiency (Duran-Trio et al., 2022) and in hippocampal neurons in a rat model of type C hepatic encephalopathy (Mosso et al., 2024). Hence, OCTOPUS can inform future research on dMRS biomarkers of pathological alterations of tissue microstructure.

Current dMRI research increasingly relies on realistic representations of tissue microstructure to improve the interpretation and modeling of diffusion signals (Abdollahzadeh et al., 2025; Chakwizira et al., 2025; Şimşek et al., 2025). Therefore, tools such as OCTOPUS, which facilitate the creation of these realistic representations, need to integrate seamlessly with existing state-of-the-art data analysis pipelines. By being fully open-source and providing a Python interface through OCTOpool, OCTOPUS is designed to be readily accessible and easily incorporated into existing research workflows. Moreover, an increasing number of Monte Carlo simulators directly support substrates described by overlapping spheres (Cottaar et al., 2026; Rafael-Patino et al., 2020; Nguyen-Duc et al., 2026a). For compatibility with other simulators, OCTOpool allows generating meshes from created cells. By avoiding dependencies on proprietary platforms or commercial licenses, OCTOPUS promotes transparent and reproducible research while facilitating community-driven development and the rapid implementation of new features.

### Limitations

The main limitation of OCTOPUS is that it currently supports the growth of individual cells only, making it readily suitable for dMRS simulations but not designed to generate voxels populated with multiple cells. This limits Monte Carlo simulations to the intracellular space and is not suited for studying phenomena where a realistic extracellular space is required, such as water dMRI and membrane exchange. However, combined with frameworks that resolve between-cell collisions such as CACTUS (Villarreal-Haro et al., 2023), we anticipate that OCTOPUS will contribute to generating densely packed gray matter substrates, also allowing a realistic representation of the extracellular space. Such substrates will allow validation of state-of-theart dMRI models for gray matter, such as SANDI (Palombo et al., 2020) and NEXI (Jelescu et al., 2022), when applied to realistic tissue geometries. Moreover, they may help to pave the way to employing more complete models of gray matter diffusion, such as SANDIX (Olesen et al., 2022), or to develop new numerical models for gray matter that account for complex features with no analytical formulation.

In its current version, the free growth approach is not capable of fully and automatically reproducing highly anisotropic cells such as pyramidal or Purkinje cells. This limitation is particularly evident in the automated cell reproduction workflow, where morphological parameters are extracted from histologically segmented cells and used directly as input in OCTOPUS. Although manually defining distinct process populations and tuning the corresponding configuration parameters can improve the resemblance of the generated morphologies to the originals, this requires user intervention. Further development of the free growth approach will be the focus of future work, for instance by applying advanced parametrization of cell branching (Aird-Rossiter et al., 2026a; Palombo et al., 2019). Nevertheless, the skeleton-based approach can already reproduce most cells accurately.

Moreover, we have not yet implemented biology-informed cell growth. Past research has characterized branching rules (Cuntz et al., 2010) and branch radius behavior after branchings (Liao et al., 2021). By implementing these rules in a future OCTOPUS release, we aim to increase the biological plausibility of the generated cells.

We observed small but consistent differences between the dMRI signals from original histological cell reconstructions and OCTOPUS-generated cells, which stem from imperfect reproduction. One simplification in OCTOPUS is representing the soma as a single sphere, which we want to extend to a more complex representation of the soma as overlapping spheres in future work.

## 5. Conclusions

OCTOPUS offers a simple yet comprehensive framework to generate biologically plausible cells with tunable morphology. This framework allows for investigating how cell geometry impacts the diffusion-weighted MRI and MRS signals. OCTOPUS can extract morphological parameter distributions from histological cell reconstructions and then reproduce these cells with high fidelity. It can also introduce morphological alterations — for instance, due to functional or pathological processes — into the reproduced cells. In combination with cell packing approaches, we envision that our work will contribute to a freely tunable generator of synthetic gray matter tissue.

## CRediT authorship contribution statement

**Malte Brammerloh:** Conceptualization, Methodology, Software, Validation, Formal analysis, Investigation, Data curation, Writing – original draft, Writing – review & editing, Visualization. **Inès de Riedmatten:** Conceptualization, Methodology, Software, Validation, Formal analysis, Investigation, Data curation, Writing – original draft, Writing – review & editing, Visualization. **Juliette Beaubis:** Methodology, Software, Validation, Writing – review & editing. **Jasmine Nguyen-Duc:** Methodology, Writing – review & editing. **Rita Oliveira:** Methodology, Writing – review & editing. **Andrés Le Boeuf Fló:** Methodology, Validation, Writing – review & editing. **Elda Fischi-Gomez:** Methodology, Writing – review & editing, Funding acquisition. **Jonathan Rafael Patiño:** Methodology, Writing – review & editing, Funding acquisition. **Ileana Jelescu:** Conceptualization, Methodology, Writing – review & editing, Supervision, Funding acquisition.

## Declaration of competing interest

The authors declare that they have no known competing financial interests or personal relationships that could have appeared to influence the work reported in this paper.

## Data availability

The OCTOPUS and OCTOpool source code will be openly available upon publication.

## Acknowledgments

We are grateful to Valerij G. Kiselev for testing our generation of cells with protrusions in other simulation environments.

This work was supported by the Swiss National Science Foundation (Proj #10.000.465 and Eccellenza #194260) and the Swiss Secretariat for Research and Innovation (ERC Starting Grant award ‘FIREPATH’ MB22.00032).

## Declaration of Generative AI and AI-assisted technologies in the writing process

During the preparation of this work, the authors used Claude (Anthropic) and ChatGPT (OpenAI) to reformulate and clarify text. After using these tools, the authors reviewed and edited the content as needed and take full responsibility for the content of the publication.

## Supplementary Materials and Methods

### OCTOpool cell geometry assessment

OCTOpool enables to create replica of histological cell reconstructions. Here, we describe the methods used to quantify the geometrical features. For a more extensive description, we refer to the OCTOpool github repository.

#### Segment lengths

A gamma distribution is fitted to the distribution of segment lengths using SciPy (Virtanen et al., 2020), and scale and shape parameters are returned. The segment length mean is calculated as the product of both.

#### Tapering

Tapering parameters were estimated by first reconstructing all unique processes by combining sequential segments. To each unique process radius, an exponential decay was fitted, mirroring the tapering model in OCTOPUS. Fits were performed in log-space for more reliable convergence. Tapering parameter plausibility is assessed by ensuring a realistic tapering length, an initial radius larger than the final radius, and positive radii. Implausible fits are excluded from the analysis, and if more than half of the tapering fits are invalid, tapering is rejected. If enough valid tapering fits are obtained, all tapering parameters are then calculated by averaging over the fitting results. If the tapering length exceeded the sum of the mean segment length and its SD, tapering is classified as continuing in secondary dendrites.

#### Beading

To estimate the beading parameters, we fit the lognormal OU model to the traced branch radii. For each branch, the radius with tapering subtracted is sampled along the branch together with the corresponding path length increments Δx. Branches shorter than a minimum length were excluded. The discretized OU equation,

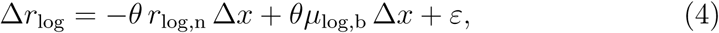

was solved by linear regression across all retained branches, with *θ* = 1/*l*_b_ shared across branches but a branch-specific mean term included for each. To accommodate large numbers of branches, the regression design matrix was constructed as a sparse indicator matrix. The correlation length was obtained as *l*_b_ = 1/*θ̂*, *µ*_log,b_ as the average of the per-branch sample means of *r*_log_, and *σ*_log,b_ from the regression residuals via 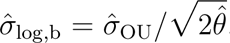, where *σ̂*_OU_ is the residual diffusion estimate. Corresponding real-space parameters (mean radius, standard deviation, and coefficient of variation) were computed analytically from the fitted lognormal parameters. If the fit was degenerate (non-finite or non-positive *θ̂*) or yielded implausible parameter values, the fitted cell was instead characterized by the empirical mean and standard deviation of r_log_ across all valid samples, and beading was not considered reliably estimable for that cell.

#### Undulation

To estimate *σ*_u_ and *l*_u_ from a cell, we fit the angular OU model to branch tangent directions. For each branch, tangents t⃗ were computed, and a per-branch preferred direction *t⃗*_0_ was estimated as the end-to-end branch direction. Segments with non-finite or zero-length arc-length increments were excluded, as were branches shorter than a minimum length or with fewer than two valid tangent increments.

The discretized relaxation term was constructed from the small-angle geodesic approximation of the model equation, Δt⃗*≈ θ ϕ*/ sin *ϕ* (*t⃗*_0_-cos *ϕ* t⃗) Δ*x*, where *ϕ* = arccos(*t⃗ ·t⃗*_0_) and *θ* = 1/*l*_u_ is shared across all branches. Stacking all tangent increments and their corresponding relaxation terms across branches, *θ* was estimated by linear regression of Δ*t⃗* against this relaxation term. The correlation length was obtained as l_u_ = 1/*θ*. The noise amplitude σ_u_ was estimated from the tangent-plane component of the regression residuals (i.e., after removing the component parallel to *t⃗*), normalized by 2Δ*x*, consistent with the two degrees of freedom of Brownian motion on the sphere.

A cell was classified as exhibiting undulation only if the fitted noise amplitude was positive and finite, the restoring rate was positive, and the resulting correlation length was finite and fell within a plausible range; otherwise, undulation was considered not detected for that cell.

### Validating OCTOpool cell geometry estimators

To validate the proposed estimators for tapering, undulation, and beading, we generated idealized test substrates for each case and examined whether the estimators could recover the ground truth. For each case, we generated elongated dendrites with a length of 1.5 mm and an average radius of 1 µm, with all possible combinations of the parameters specified below. Then we plotted the ground truth parameters against the estimated values.

#### Tapering

We generated tapered dendrites with tapering lengths of 1, 5, 10, 25, 50 µm, and tapering start radii of 1, 2, 3, 4, 5 µm, where the first start radius implies no tapering. The tapering estimator can retrieve almost all parameters with high accuracy, except for cases of short tapering lengths and high tapering start radii, where too few points are available for an accurate fit (Fig. S1).

#### Undulation

We created undulated dendrites with *σ_u_* = 0.1, 0.2, 0.3, 0.4, and correlation lengths of 2, 5, 10, 15 µm. The undulation estimator retrieved σ*_u_* with high accuracy in all cases, except for the highest *σ_u_*, where *σ_u_* was slightly underestimated (Fig. S2). The correlation length estimation was less precise for increasing correlation lengths, but showed no systematic bias (Fig. S2).

**Figure S1:**
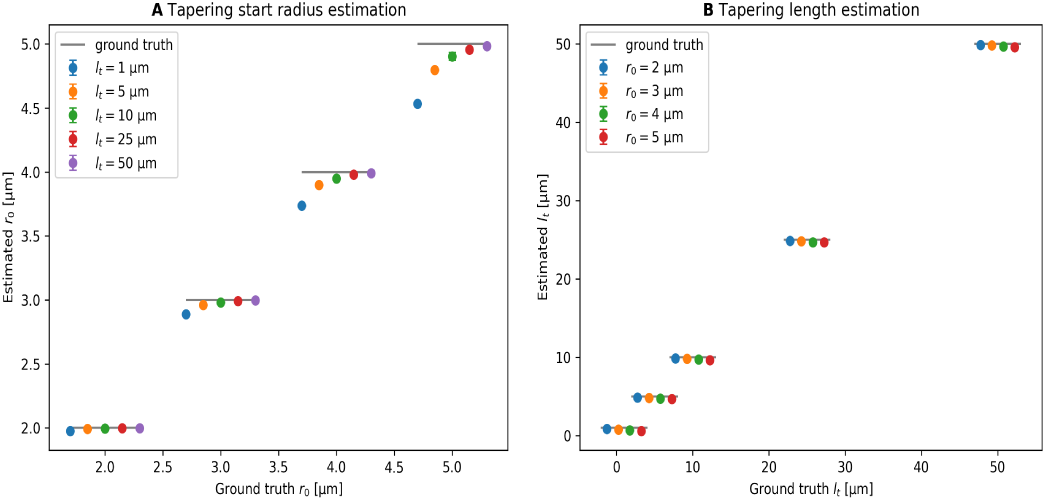
Evaluation of the tapering estimator. The horizontal gray bars indicate the ground truth value.

**Figure S2:**
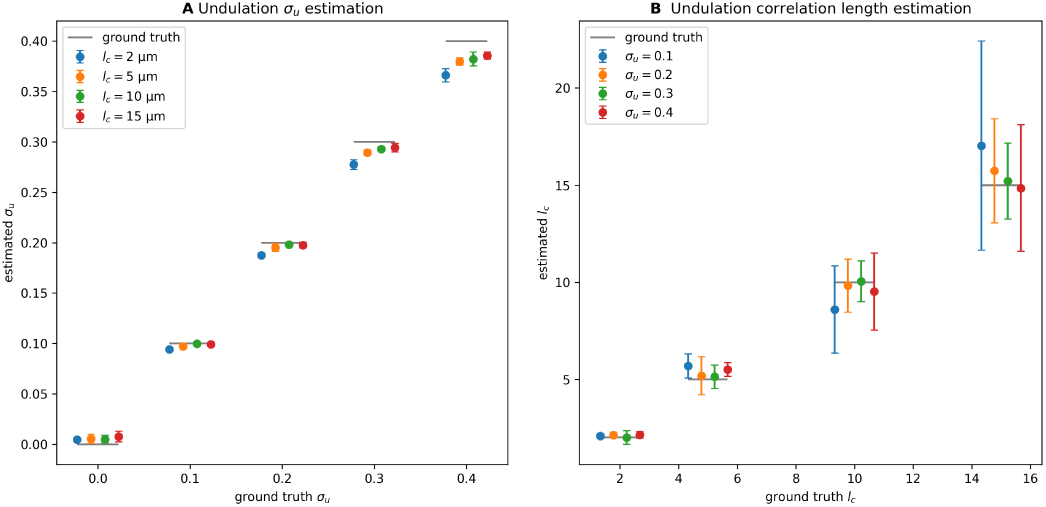
Evaluation of the undulation estimator. The horizontal gray bars indicate the ground truth value.

**Figure S3:**
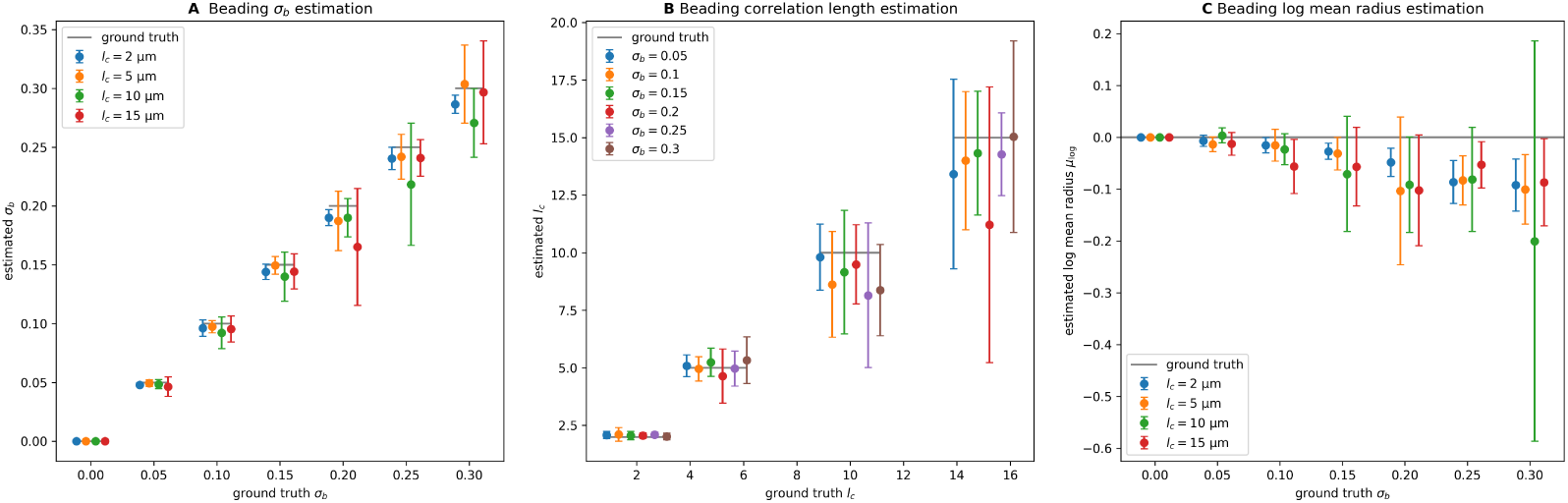
Evaluation of the beading estimator. The horizontal gray bars indicate the ground truth value.

#### Beading

We created beaded dendrites with σ*_b_* = 0, 0.05, 0.1, . . ., 0.3 µm and correlation lengths of 2, 5, 10, 15 µm. The parameter σ*_b_* is estimated with high accuracy in all cases (Fig. S3A). The accuracy of the correlation length is overall lower and has a slight negative bias for strong beading (Fig. S3B). Both accuracies decrease with increasing σ*_b_* (Fig. S3A,B). We noticed that stronger beading leads to reduced accuracy and a slight negative bias in the log-mean radius estimate (Fig. S3C).

---

**Table S1:**
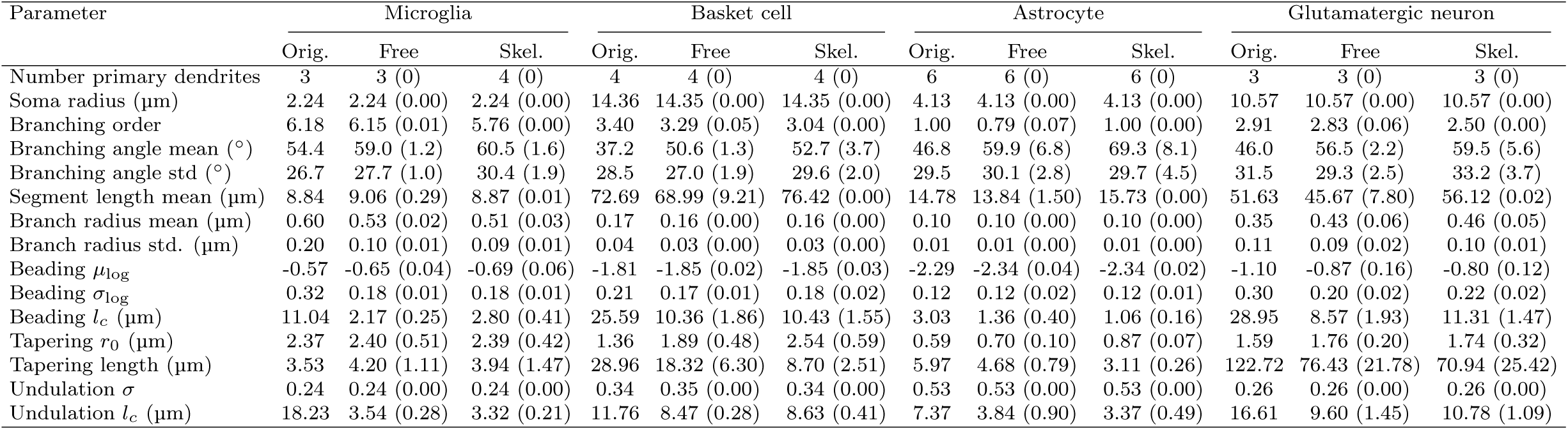
Cell reproduction analysis I.

**Table S2:**
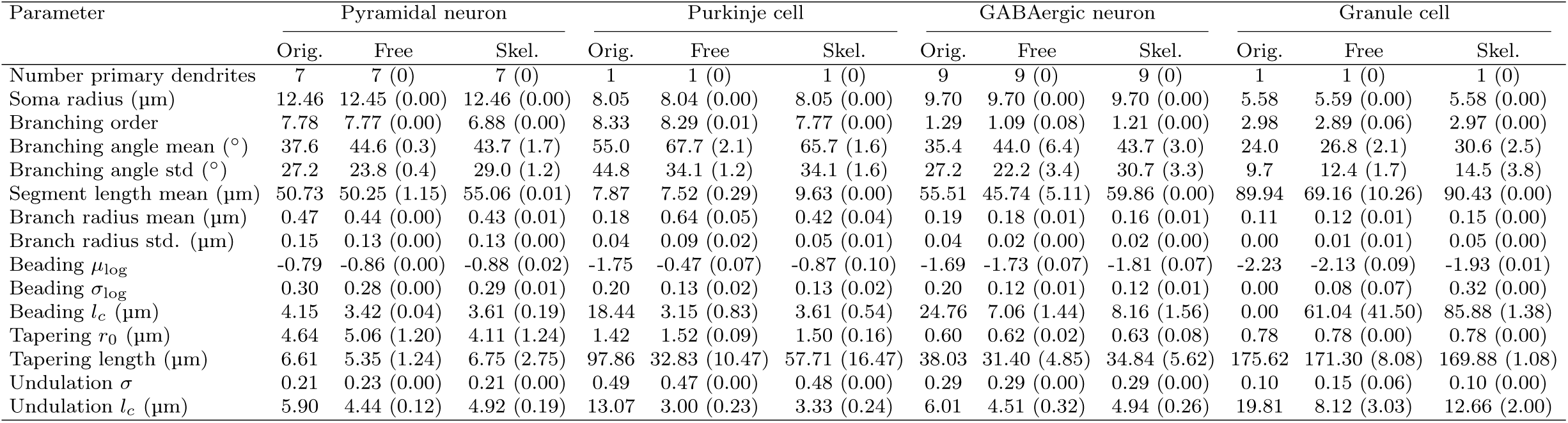
Cell reproduction analysis II.

## Notes

### Competing Interest Statement

The authors have declared no competing interest.

## References

Abdollahzadeh, A., Coronado-Leija, R., Lee, H.H., Sierra, A., Fieremans, E., Novikov, D.S., 2025. Scattering approach to diffusion quantifies axonal damage in brain injury. Nature Communications 16, 9808. URL: https://www.nature.com/articles/s41467-025-64793-1, doi:10.1038/s41467-025-64793-1. publisher: Nature Publishing Group.

Aird-Rossiter, C., Zhang, H., Alexander, D.C., Jones, D.K., Palombo, M., 2026a. Decoding gray matter, large-scale analysis of brain cell morphometry to inform microstructural modeling of diffusion MR signals. Communications Biology 9, 138. URL: https://www.nature.com/articles/s42003-025-09353-5, doi:10.1038/s42003-025-09353-5.

Aird-Rossiter, C., Şimşek, K., Jallais, M., Jones, D.K., Kanari, L., Palombo, M., 2026b. Contextual Cellular Growth (ConCeG) of neural cells for realistic grey matter tissue generation for diffusion MRI simulations. URL: http://arxiv.org/abs/2607.03286, doi:10.48550/arXiv.2607.03286.

Alexander, D.C., Dyrby, T.B., Nilsson, M., Zhang, H., 2019. Imaging brain microstructure with diffusion MRI: practicality and applications. NMR in Biomedicine 32, e3841. URL: https://analyticalsciencejournals.onlinelibrary.wiley.com/doi/10.1002/nbm.3841, doi:10.1002/nbm.3841.

Ascoli, G.A., Donohue, D.E., Halavi, M., 2007. NeuroMorpho.Org: A Central Resource for Neuronal Morphologies. Journal of Neuroscience 27, 9247– 9251. URL: https://www.jneurosci.org/content/27/35/9247, doi:10.1523/JNEUROSCI.2055-07.2007.

Augustin, E., Vinasco-Sandoval, T., Riquelme-Perez, M., Plassard, D., Gaudin, M., Aurégan, G., Mitja, J., Bernier, S., Joséphine, C., Petit, F., Jan, C., Hérard, A.S., Gaillard, M.C., Launay, A., Faivre, E., Buée, L., Boutillier, A.L., Blum, D., Bemelmans, A.P., Bonvento, G., Cambon, K., 2025. Hippocampal Astrocyte Morphology Follows an Unexpected Trajectory With Age in a Transgenic Rodent Model of Tauopathy. Glia 73, 1502–1519. URL: https://onlinelibrary.wiley.com/doi/abs/10.1002/glia.70019, doi:10.1002/glia.70019. _eprint: https://onlinelibrary.wiley.com/doi/pdf/10.1002/glia.70019.

Brammerloh, M., Morawski, M., Friedrich, I., Reinert, T., Lange, C., Pelicon, P., Vavpetič, P., Jankuhn, S., Jäger, C., Alkemade, A., Balesar, R., Pine, K., Gavriilidis, F., Trampel, R., Reimer, E., Arendt, T., Weiskopf, N., Kirilina, E., 2021. Measuring the iron content of dopaminergic neurons in substantia nigra with MRI relaxometry. NeuroImage 239, 118255. URL: https://linkinghub.elsevier.com/retrieve/pii/S1053811921005322, doi:10.1016/j.neuroimage.2021.118255.

Budde, M.D., Frank, J.A., 2010. Neurite beading is sufficient to decrease the apparent diffusion coefficient after ischemic stroke. Proceedings of the National Academy of Sciences of the United States of America 107, 14472– 14477. doi:10.1073/pnas.1004841107.

Callaghan, P., Jolley, K., Lelievre, J., 1979. Diffusion of water in the endosperm tissue of wheat grains as studied by pulsed field gradient nuclear magnetic resonance. Biophysical Journal 28, 133–141. URL: https://linkinghub.elsevier.com/retrieve/pii/S0006349579851644, doi:10.1016/S0006-3495(79)85164-4.

Callaghan, R., Alexander, D.C., Palombo, M., Zhang, H., 2020. ConFiG: Contextual Fibre Growth to generate realistic axonal packing for diffusion MRI simulation. NeuroImage 220, 117107. URL: https://www.sciencedirect.com/science/article/pii/S1053811920305930, doi:10.1016/j.neuroimage.2020.117107.

Chakwizira, A., Şimşek, K., Szczepankiewicz, F., Palombo, M., Nilsson, M., 2025. The role of dendritic spines in water exchange measurements with diffusion MRI: Double Diffusion Encoding and free-waveform MRI. URL: http://arxiv.org/abs/2504.21537, doi:10.48550/arXiv.2504.21537. arXiv:2504.21537 [physics].

Cottaar, M., Zheng, Z., Miller, K.L., Tendler, B.C., Jbabdi, S., 2026. Multimodal Monte Carlo MRI simulator of tissue microstructure. Imaging Neuroscience 4, IMAG.a.1177. URL: https://direct.mit.edu/imag/article/doi/10.1162/IMAG.a.1177/135558/Multi-modal-Monte-Carlo-MRI-simulator-of-tissue, doi:10.1162/IMAG.a.1177.

Cuntz, H., Forstner, F., Borst, A., Häusser, M., 2010. One Rule to Grow Them All: A General Theory of Neuronal Branching and Its Practical Application. PLOS Computational Biology 6, e1000877. URL: https://journals.plos.org/ploscompbiol/article?id=10.1371/journal.pcbi.1000877, doi:10.1371/journal.pcbi.1000877. publisher: Public Library of Science.

Duran-Trio, L., Fernandes-Pires, G., Grosse, J., Soro-Arnaiz, I., RouxPetronelli, C., Binz, P.A., De Bock, K., Cudalbu, C., Sandi, C., Braissant, O., 2022. Creatine transporter–deficient rat model shows motor dysfunction, cerebellar alterations, and muscle creatine deficiency without muscle atrophy. Journal of Inherited Metabolic Disease 45, 278–291. URL: https://onlinelibrary.wiley.com/doi/abs/10.1002/jimd.12470, doi:10.1002/jimd.12470. _eprint: https://onlinelibrary.wiley.com/doi/pdf/10.1002/jimd.12470.

Fieremans, E., Lee, H.H., 2018. Physical and numerical phantoms for the validation of brain microstructural MRI: A cookbook. NeuroImage 182, 39–61. doi:10.1016/j.neuroimage.2018.06.046.

Ginsburger, K., Matuschke, F., Poupon, F., Mangin, J.F., Axer, M., Poupon, C., 2019. MEDUSA: A GPU-based tool to create realistic phantoms of the brain microstructure using tiny spheres. NeuroImage 193, 10–24. URL: https://www.sciencedirect.com/science/article/pii/S105381191930151X, doi:10.1016/j.neuroimage.2019.02.055.

Ianus, A., Alexander, D.C., Zhang, H., Palombo, M., 2021. Mapping complex cell morphology in the grey matter with double diffusion encoding MR: A simulation study. NeuroImage 241, 118424. URL: https://www.sciencedirect.com/science/article/pii/S1053811921006996, doi:10.1016/j.neuroimage.2021.118424.

Jelescu, I.O., Palombo, M., Bagnato, F., Schilling, K.G., 2020. Challenges for biophysical modeling of microstructure. Journal of Neuroscience Methods 344, 108861. doi:10.1016/j.jneumeth.2020.108861.

Jelescu, I.O., de Skowronski, A., Geffroy, F., Palombo, M., Novikov, D.S., 2022. Neurite Exchange Imaging (NEXI): A minimal model of diffusion in gray matter with inter-compartment water exchange. NeuroImage 256, 119277. URL: https://www.sciencedirect.com/science/article/pii/S1053811922003986, doi:10.1016/j.neuroimage.2022.119277.

Lee, H.H., Novikov, D.S., Fieremans, E., Huang, S.Y., 2025. Revealing membrane integrity and cell size from diffusion kurtosis time dependence. Magnetic Resonance in Medicine 93, 1329–1347. URL: https://onlinelibrary.wiley.com/doi/abs/10.1002/mrm.30335, doi:10.1002/mrm.30335. _eprint: https://onlinelibrary.wiley.com/doi/pdf/10.1002/mrm.30335.

Lee, H.H., Papaioannou, A., Kim, S.L., Novikov, D.S., Fieremans, E., 2020a. A time-dependent diffusion MRI signature of axon caliber variations and beading. Communications biology 3, 354. URL: https://www.nature.com/articles/s42003-020-1050-x. publisher: Nature Publishing Group UK London.

Lee, H.H., Papaioannou, A., Novikov, D.S., Fieremans, E., 2020b. In vivo observation and biophysical interpretation of time-dependent diffusion in human cortical gray matter. NeuroImage 222, 117054. URL: https:// www.sciencedirect.com/science/article/pii/S1053811920305401, doi:10.1016/j.neuroimage.2020.117054.

Liao, M., Liang, X., Howard, J., 2021. The narrowing of dendrite branches across nodes follows a well-defined scaling law. Proceedings of the National Academy of Sciences 118, e2022395118. URL: https://www.pnas.org/doi/10.1073/pnas.2022395118, doi:10.1073/pnas.2022395118. publisher: Proceedings of the National Academy of Sciences.

Ligneul, C., Najac, C., Döring, A., Beaulieu, C., Branzoli, F., Clarke, W.T., Cudalbu, C., Genovese, G., Jbabdi, S., Jelescu, I., Karampinos, D., Kreis, R., Lundell, H., Marjańska, M., Möller, H.E., Mosso, J., Mougel, E., Posse, S., Ruschke, S., Simsek, K., Szczepankiewicz, F., Tal, A., Tax, C., Oeltzschner, G., Palombo, M., Ronen, I., Valette, J., 2024. Diffusion-weighted <span style=“font-variant:small-caps;”>MR</SPAN> spectroscopy: Consensus, recommendations, and resources from acquisition to modeling. Magnetic Resonance in Medicine 91, 860–885. URL: https://onlinelibrary.wiley.com/doi/10.1002/mrm.29877, doi:10.1002/mrm.29877.

Ligneul, C., Qiu, L., Clarke, W.T., Jbabdi, S., Palombo, M., Lerch, J.P., 2025. Diffusion MRS tracks distinct trajectories of neuronal development in the cerebellum and thalamus of rat neonates. eLife 13. URL: https://elifesciences.org/reviewed-preprints/96625, doi:10.7554/eLife.96625.3. publisher: eLife Sciences Publications Limited.

Mosso, J., Briand, G., Pierzchala, K., Simicic, D., Sierra, A., Abdollahzadeh, A., Jelescu, I.O., Cudalbu, C., 2024. Diffusion of brain metabolites highlights altered brain microstructure in type C hepatic encephalopathy: a 9.4 T preliminary study. Frontiers in Neuroscience 18, 1344076. URL: https://www.frontiersin.org/articles/10.3389/fnins.2024.1344076/full, doi:10.3389/fnins.2024.1344076.

Nguyen-Duc, J., Brammerloh, M., Cherchali, M., De Riedmatten, I., Pérot, J.B., Rafael-Patiño, J., Jelescu, I.O., 2026a. CATERPillar: a flexible framework for generating white matter numerical substrates with incorporated glial cells. Medical Image Analysis 110, 103946. URL: https://www.sciencedirect.com/science/article/pii/S1361841526000150, doi:10.1016/j.media.2026.103946.

Nguyen-Duc, J., Uhl, Q., Oliveira, R., Rafael-Patiño, J., Jelescu, I.O., 2026b. Validating the Standard Model of diffusion MRI in white matter with Numerical Substrates. URL: https://www.biorxiv.org/content/10.64898/2026.01.28.702302v1, doi:10.64898/2026.01.28.702302. iSSN: 2692-8205 Pages: 2026.01.28.702302 Section: New Results.

Novikov, D.S., Fieremans, E., Jespersen, S.N., Kiselev, V.G., 2019. Quantifying brain microstructure with diffusion MRI: Theory and parameter estimation. NMR in Biomedicine 32, e3998. URL: https://onlinelibrary.wiley.com/doi/abs/10.1002/nbm.3998, doi:10.1002/nbm.3998. _eprint: https://onlinelibrary.wiley.com/doi/pdf/10.1002/nbm.3998.

Novikov, D.S., Jensen, J.H., Helpern, J.A., Fieremans, E., 2014. Revealing mesoscopic structural universality with diffusion. Proceedings of the National Academy of Sciences 111, 5088–5093. URL: https://www.pnas.org/doi/abs/10.1073/pnas.1316944111, doi:10.1073/pnas.1316944111. _eprint: https://www.pnas.org/doi/pdf/10.1073/pnas.1316944111.

Ofer, N., Berger, D.R., Kasthuri, N., Lichtman, J.W., Yuste, R., 2021. Ultrastructural analysis of dendritic spine necks reveals a continuum of spine morphologies. Developmental neurobiology 81, 746–757. URL: https://www.ncbi.nlm.nih.gov/pmc/articles/PMC8852350/, doi:10.1002/dneu.22829.

Olesen, J.L., Østergaard, L., Shemesh, N., Jespersen, S.N., 2022. Diffusion time dependence, power-law scaling, and exchange in gray matter. NeuroImage 251, 118976. URL: https://www.sciencedirect.com/science/article/pii/S1053811922001057, 10.1016/j.neuroimage.2022.118976.

Oliveira, R., Nguyen-Duc, J., Brammerloh, M., Jelescu, I., 2026. Validating Neurite EXchange Imaging (NEXI) using diffusion Monte Carlo simulations in realistic numerical gray matter substrates. URL: https://www.biorxiv.org/content/10.64898/2026.02.11.705314v1, doi:10.64898/2026.02.11.705314. iSSN: 2692-8205 Pages: 2026.02.11.705314 Section: New Results.

Palombo, M., Alexander, D.C., Zhang, H., 2019. A generative model of realistic brain cells with application to numerical simulation of the diffusion-weighted MR signal. NeuroImage 188, 391–402. URL: https://www.sciencedirect.com/science/article/pii/S1053811918321694, doi:10.1016/j.neuroimage.2018.12.025.

Palombo, M., Ianus, A., Guerreri, M., Nunes, D., Alexander, D.C., Shemesh, N., Zhang, H., 2020. SANDI: A compartment-based model for non-invasive apparent soma and neurite imaging by diffusion MRI. NeuroImage 215, 116835. URL: https://www.sciencedirect.com/science/article/pii/S1053811920303220, doi:10.1016/j.neuroimage.2020.116835.

Palombo, M., Ligneul, C., Hernandez-Garzon, E., Valette, J., 2017. Can we detect the effect of spines and leaflets on the diffusion of brain intracellular metabolites? NeuroImage 182, 283–293. doi:10.1016/j.neuroimage.2017.05.003.

Palombo, M., Ligneul, C., Najac, C., Le Douce, J., Flament, J., Escartin, C., Hantraye, P., Brouillet, E., Bonvento, G., Valette, J., 2016. New paradigm to assess brain cell morphology by diffusion-weighted MR spectroscopy in vivo. Proceedings of the National Academy of Sciences 113, 6671–6676. URL: https://www.pnas.org/doi/abs/10.1073/pnas.1504327113, doi:10.1073/pnas.1504327113. publisher: Proceedings of the National Academy of Sciences.

Panagiotaki, E., Walker-Samuel, S., Siow, B., Johnson, S.P., Rajkumar, V., Pedley, R.B., Lythgoe, M.F., Alexander, D.C., 2014. Noninvasive quantification of solid tumor microstructure using VERDICT MRI. Cancer Research 74, 1902–1912. doi:10.1158/0008-5472.CAN-13-2511.

Rafael-Patino, J., Romascano, D., Ramirez-Manzanares, A., CanalesRodríguez, E.J., Girard, G., Thiran, J.P., 2020. Robust Monte-Carlo Simulations in Diffusion-MRI: Effect of the Substrate Complexity and Parameter Choice on the Reproducibility of Results. Frontiers in Neuroinformatics 14. URL: https://www.frontiersin.org/journals/neuroinformatics/articles/10.3389/fninf.2020.00008/full, doi:10.3389/fninf.2020.00008. publisher: Frontiers.

Spencer, A.P.C., Nguyen-Duc, J., de Riedmatten, I., Szczepankiewicz, F., Jelescu, I.O., 2025. Mapping grey and white matter activity in the human brain with isotropic ADC-fMRI. Nature Communications 16, 5036. URL: https://www.nature.com/articles/s41467-025-60357-5, doi:10.1038/s41467-025-60357-5.

Stanisz, G.J., Szafer, A., Wright, G.A., Henkelman, R.M., 1997. An analytical model of restricted diffusion in bovine optic nerve. Magnetic Resonance in Medicine 37, 103–111. doi:10.1002/mrm.1910370115.

Tecuatl, C., Ljungquist, B., Ascoli, G.A., 2024. Accelerating the continuous community sharing of digital neuromorphology data. FASEB BioAdvances 6, 207–221. URL: https://pmc.ncbi.nlm.nih.gov/articles/PMC11226999/, doi:10.1096/fba.2024-00048.

Uhlenbeck, G.E., Ornstein, L.S., 1930. On the Theory of the Brownian Motion. Physical Review 36, 823–841. URL: https://link.aps.org/doi/10.1103/PhysRev.36.823, doi:10.1103/PhysRev.36.823. publisher: American Physical Society.

Veraart, J., Fieremans, E., Novikov, D.S., 2019. On the scaling behavior of water diffusion in human brain white matter. NeuroImage 185, 379–387. URL: https://www.sciencedirect.com/science/article/pii/S1053811918319475, 10.1016/j.neuroimage.2018.09.075.

Villarreal-Haro, J.L., Gardier, R., Canales-Rodríguez, E.J., Fischi-Gomez, E., Girard, G., Thiran, J.P., Rafael-Patiño, J., 2023. CACTUS: a computational framework for generating realistic white matter microstructure substrates. Frontiers in Neuroinformatics 17. URL: https://www.frontiersin.org/journals/neuroinformatics/articles/10.3389/fninf.2023.1208073/full, doi:10.3389/fninf.2023.1208073. publisher: Frontiers.

Virtanen, P., Gommers, R., Oliphant, T.E., Haberland, M., Reddy, T., Cournapeau, D., Burovski, E., Peterson, P., Weckesser, W., Bright, J., van der Walt, S.J., Brett, M., Wilson, J., Millman, K.J., Mayorov, N., Nelson, A.R.J., Jones, E., Kern, R., Larson, E., Carey, C.J., Polat, İ., Feng, Y., Moore, E.W., VanderPlas, J., Laxalde, D., Perktold, J., Cimrman, R., Henriksen, I., Quintero, E.A., Harris, C.R., Archibald, A.M., Ribeiro, A.H., Pedregosa, F., van Mulbregt, P., 2020. SciPy 1.0: fundamental algorithms for scientific computing in Python. Nature Methods 17, 261–272. URL: https://www.nature.com/articles/s41592-019-0686-2, doi:10.1038/s41592-019-0686-2. number: 3.

Voronova, A.K., Grigoriou, A., Bernatowicz, K., Simonetti, S., Serna, G., Roson, N., Escobar, M., Vieito, M., Nuciforo, P., Toledo, R., Garralda, E., Fieremans, E., Novikov, D.S., Palombo, M., Perez-Lopez, R., Grussu, F., 2025. SpinFlowSim: A blood flow simulation framework for histology-informed diffusion MRI microvasculature mapping in cancer. Medical Image Analysis 102, 103531. URL: https://www.sciencedirect.com/science/article/pii/S1361841525000799, doi:10.1016/j.media.2025.103531.

Weiskopf, N., Edwards, L.J., Helms, G., Mohammadi, S., Kirilina, E., 2021. Quantitative magnetic resonance imaging of brain anatomy and in vivo histology. Nature Reviews Physics, 1–19URL: https://www.nature.com/articles/s42254-021-00326-1, doi:10.1038/s42254-021-00326-1. bandiera_abtest: a Cg_type: Nature Research Journals Primary_atype: Reviews Publisher: Nature Publishing Group Subject_term: Biological physics;Biophysics;Imaging techniques Subject_term_id: biologicalphysics;biophysics;imaging-techniques.

Winther, S., Peulicke, O., Andersson, M., Kjer, H.M., Bærentzen, J.A., Dyrby, T.B., 2024. Exploring white matter dynamics and morphology through interactive numerical phantoms: the White Matter Generator. Frontiers in Neuroinformatics 18. URL: https://www.frontiersin.org/journals/neuroinformatics/articles/10.3389/fninf.2024.1354708/full, doi:10.3389/fninf.2024.1354708. publisher: Frontiers.

Zhang, H., Schneider, T., Wheeler-Kingshott, C.A., Alexander, D.C., 2012. NODDI: Practical *in vivo* neurite orientation dispersion and density imaging of the human brain. NeuroImage 61, 1000–1016. URL: https://www.sciencedirect.com/science/article/pii/S1053811912003539, doi:10.1016/j.neuroimage.2012.03.072.

Şimşek, K., Chakwizira, A., Nilsson, M., Palombo, M., 2025. The role of dendritic spines in water exchange measurements with diffusion MRI: Time-Dependent Single Diffusion Encoding MRI. URL: http://arxiv.org/abs/2506.18229, doi:10.48550/arXiv.2506.18229. arXiv:2506.18229[physics].

